# The Type II Secretion System Utilizes AsmA-like Protein GspN to Facilitate Transport of Lipoproteins to the Cell Surface in *Acinetobacter baumannii*

**DOI:** 10.1101/2025.09.09.675180

**Authors:** Cameron S. Roberts, Colby Gura, Maria Sandkvist

## Abstract

Gram-negative bacteria employ the Type II Secretion System (T2SS) to not only secrete an array of soluble effectors such as toxins to the extracellular space, but also to facilitate the surface localization of enzymes and adhesins that are beneficial to life in different environments. For example, the pullulan degrading enzyme pullulanase (PulA) from *Klebsiella pneumoniae* and the recently discovered adhesin InvL from *Acinetobacter baumannii* are initially expressed with a lipobox containing signal peptide, resulting in their N-terminal acylation and subsequent surface anchoring after T2SS mediated export. While outer membrane translocation of both soluble and surface associated T2SS effectors depends on the T2SS secretin GspD, it is unclear how lipoproteins are accommodated by the T2SS during transport to the cell surface. Here, we identify a role for GspN in the outer membrane translocation of InvL in the opportunistic pathogen *A. baumannii*. Additional putative lipoproteins are found to have a similar GspN dependence for secretion, while soluble proteins are secreted independently of GspN. We demonstrate that a specific sorting motif C-terminal to the lipobox is required for GspN-dependent surface localization. Based on structural predictions, GspN belongs to the larger AsmA-like protein family that includes both eukaryotic and prokaryotic members. This protein family has been implicated in phospholipid transport, but here we expand the role for this family to include transport of lipoproteins. We also confirm that the GspN homolog PulN is required for PulA surface localization in *K. pneumoniae*.

**Significance Statement:** The Type II Secretion (T2SS) is considered a virulence factor of gram-negative pathogens such as *A. baumannii*. The role of one of the components of the T2SS, GspN, has been unclear and many studies across multiple model systems have reported that GspN is dispensable for protein secretion. Here, we characterize the transport of a subset of proteins t by the T2SS and show that GspN is required for their outer membrane translocation and surface localization. GspN belongs to the AsmA-like protein family, which to date has only been implicated in the transport of phospholipids. This study demonstrates that, in addition to transport of phospholipids, this class of proteins also facilitates the secretion of proteins.

## Introduction

The Type II Secretion System (T2SS) is responsible for the secretion of a variety of effector molecules to the extracellular space in select gram-negative bacteria (1-3) including proteases, lipases, carbohydrate active enzymes and toxins that support environmental persistence and host colonization (2, 4, 5). While many effector molecules are freely released to the extracellular space following outer membrane translocation, several are retained on the bacterial cell surface (1, 6). Cell surface associated effectors include PulA from *K. pneumoniae*, heat-labile enterotoxin from *Escherichia coli*, PnlH from *Dickeya dadantii*, VesB from *V. cholerae*, and InvL from *A. baumannii* (7-11). Several distinct mechanisms of surface association exist. For example, VesB is retained on the cell surface via a posttranslational modification of its C-terminus with a glycerophosphoethanolamine-containing moiety (12) and PnlH is produced with a non-cleavable TAT signal peptide, which establishes surface association (9). In the case of heat-labile enterotoxin, the B subunit of the toxin binds to lipopolysaccharides (10). Finally, PulA and InvL are produced with a signal peptide containing a lipobox, resulting in lipidation at a conserved cysteine residue and processing by the signal peptidase II prior to secretion and surface anchoring. Given their hydrophobic moiety, it is unclear how the T2SS can accommodate lipoproteins during their transit of the periplasmic space.

The T2SS complex spans the entirety of the cell envelope and is encoded by 13-16 genes often present in a single operon or *gsp* locus (2). Classically, each of these genes is considered essential for the function of the entire system with one notable exception (2, 13, 14). Despite demonstrating the presence of the T2SS gene product GspN in purified secretion complexes from *K. pneumoniae* (15), it has been found to be dispensable for the secretion of effector molecules from a variety of bacteria (2, 14). While not encoded by all bacteria expressing T2SS, in addition to *A. baumannii*, GspN is also found in the notable pathogens *V. cholerae, K. pneumoniae*, and *Burkholderia pseudomallei*. It was previously speculated that GspN may only be needed for the efficient secretion of effector molecules under specific growth conditions or only required for the secretion of a specific subset of effector molecules not yet identified (2).

Here, we probe the requirement for GspN in the secretion of a subset of T2SS effectors that are retained at the cell surface via their acylated N-terminus, such as InvL. These proteins are believed to be posttranslationally modified with two acyl groups by Lgt, before cleavage of their leader peptide by Lsp II, and finally modified with a third acyl group by Lnt based on data from *E. coli* (16). GspN is predicted to adopt a fold similar to the so called AsmA-like proteins, members of a larger protein family including prokaryotic and eukaryotic members involved in phospholipid transport (17). Bacterial AsmA-like proteins can be quite large, some large enough to span the entirety of the periplasmic space and have been linked to the transport of phospholipids between the inner and outer membranes (18). Other members such as GspN are significantly smaller and have yet to be ascribed a physiological function (19). To our knowledge a direct role in the transport of proteins has yet to be demonstrated for AsmA-like proteins. Using a contemporary clinical isolate of *A. baumannii*, we show that the known T2SS effector InvL, produced with a lipobox, requires GspN for efficient outer membrane translocation and surface localization. We have identified additional T2SS dependent lipoproteins and validated GspN dependence for several of them. Both GspN and a large number of the T2SS dependent lipoproteins are highly conserved amongst *A. baumannii* isolates. PulA from *K. pneumoniae* also relies on the GspN homolog PulN for surface localization and the T2SS sorting motif is present within lipoproteins from *B. pseudomallei* and *Shewanella oneidensis*. Overall, we describe a novel function for an AsmA-like protein and shed light on the mysterious role of GspN that had previously eluded illumination.

## Results

### *The gspN* gene is highly conserved amongst a subset of T2SS containing bacteria

The T2SS consists of 13-16 proteins that form a complex nanomachine that spans the entire cell envelope (**Figure 1A**) (1). This nanomachine is distributed across a subset of gram-negative bacteria where studies have focused on select model organisms including *E. coli, K. pneumoniae, D. dadantii, Pseudomonas aeruginosa, A. baumannii, V. cholerae, Xanthomonas campestris, Legionella pneumophila, B. pseudomallei, and S. oneidensis* (2, 9, 20-26). Of these species, the structural genes *gspC-gspM* are universal, while *gspN* is only found in *K. pneumoniae, V. cholerae, A. baumannii, B. pseudomallei, and S. oneidensis*. (**Figure 1B**). Given past speculation on the role of, or lack thereof, for GspN in T2SS mediated protein translocation, we examined the conservation of *gpsN* across three of these species. Of the fully assembled *K. pneumoniae* genomes extracted from NCBI we identified 3702 genomes with *gspN* out of 3703 total analyzed. In the single genome lacking *gspN* (CP110177.1) we identified *gspE*, but none of the other *gsp* genes, suggesting that this strain lacks a functional T2SS. Upon examination of 211 fully assembled *V. cholerae* genomes, nine genomes were found to lack all *gsp* genes including *gspN*. Of the 882 *A. baumannii* genomes, the vast majority (873) contains *gspN*. Those lacking *gspN* are also missing multiple other *gsp* genes. Previously, *A. baumannii* strain AB307-0294 (CP001172.2) has been postulated to lack *gspN*, but harbor the other genes required for T2SS function (27). However, our analysis revealed that this strain indeed does have a *gspN* copy whose primary amino acid sequence (ATY45646.1) is highly conserved with GspN from other *A. bauamannii* strains. Collectively, our findings suggest that there is evolutionary pressure for species that have a T2SS and *gspN*, to conserve said gene. We speculate that *gspN* conservation in a select few T2SS containing bacteria implies a specific role for GspN in the transport of a subset of effector proteins.

**Figure 1.**
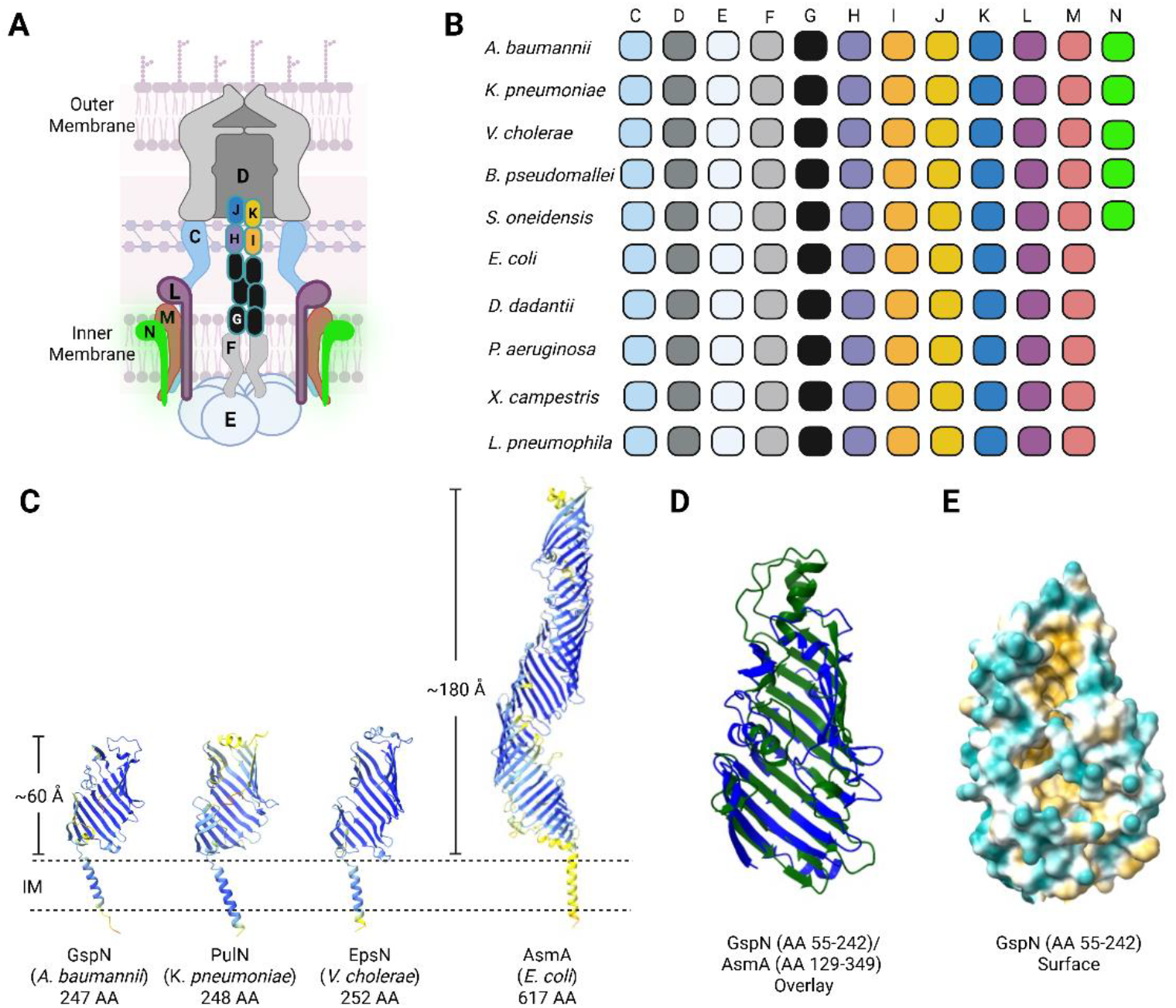
GspN is structurally homologous to AsmA-Like proteins. **A**. Model of the Type II Secretion System (T2SS) nanomachine spanning the cell envelope of gram-negative bacteria. Core components are indicated with letter nomenclature and a hypothetical position of GspN (in green) is shown. Prepilin peptidase and pilotin are not shown. **B**. Conservation of genes coding for T2SS core components from representative species. Specific strains analyzed include *gspN*-containing strains *A. baumannii* G414, *K. pneumoniae* KPPR1, *V. cholerae* N16961, *B. pseudomallei* 688, and *S. oneidensis* MR-1 and those lacking *gspN E. coli* E2348/69, *D. dadantii* 3937, *P. aeruginosa* PAO1, *X. campestris* 85-10 and *L. pneumoniae* 130b. **C**. Predicted AlphaFold3 structure of GspN (encoded by RE180_16620) from *A. baumannii* G414 was generated, while the predicted structure of PulN from *K. pneumoniae* (P15753), EpsN from *V. cholerae* (P45784), and AsmA from *E. coli* (P28249) were extracted from the AlphaFold database. Structures are colored by pLDDT confidence score in dark blue (pLDDT > 90), cyan (70 < pLDDT < 90) and yellow (50 < pLDDT < 70). Aligned residue versus expected error for GspN is shown in **Supplemental Figure 1**. Proteins are manually oriented in the inner membrane (IM). **D**. Superposition of GspN (blue) and AsmA (green) predicted structures. The predicted structures of indicated amino acid residues (AA) are shown with a RMSD of 8.4. GspN is oriented as in **C**, while AsmA has been inverted relative to its horizontal axis. **E**. Surface representation of the predicted GspN structure is colored by hydrophobicity with cyan representing most hydrophilic and gold, most hydrophobic.

### GspN is structurally homologous to bacterial AsmA-like proteins

To gain insight into the potential function of GspN, we examined the AlphaFold-predicted structures of the GspN homologs from *A. baumannii, K. pneumoniae*, and *V. cholerae* (**Figure 1C**). *A. baumannii* GspN is predicted with high confidence (**Sup. Figure 1**) to adopt a β-taco fold that is reminiscent of eukaryotic and prokaryotic proteins shown to facilitate transport of lipids (17). Both the GspN homologs PulN from *K. pneumoniae* and EpsN from *V. cholerae* are predicted to adopt a similar fold (**Figure 1C**). Consistent with previous results demonstrating that *K. pneumoniae* PulN co-purifies with the inner membrane complex of the T2SS consisting of PulC, PulL, and PulM, GspN is predicted to harbor a single alpha helix that could orient the protein in the inner membrane with the β-taco fold facing the periplasmic space. Bacterial proteins with similar orientation and β-taco folds fall into the AsmA-like protein family, named for the *E. coli* protein AsmA shown in **Figure 1C**. This family carries repeating β-taco folds that in some cases are predicted to span the entirety of the periplasmic space. When we compared the structures of GspN and AsmA, we observed significant similarity in the first β-taco fold of AsmA with the globular domain of GspN with a root-mean square deviation (RMSD) of 8.4 (**Figure 1D**), although GspN is predicted to be significantly shorter in span (60Å vs 180Å). We note that the structural similarity between GspN and AsmA requires a 180° rotation over the horizontal axis of AsmA. We also visualized the globular domain of GspN with surface representation and note the hydrophobic pocket contained within the β-taco fold (**Figure 1E**).

### *A. baumannii* strains express multiple putative T2SS effectors with a lipobox containing signal peptide

Given the structural similarity between GspN and AsmA, we hypothesized that GspN might support the transport of lipoproteins. We therefore searched for lipoproteins in three published datasets on the T2SS dependent secretomes from *A. baumannii* strains UBPA1, 17978, and AB5075 (11, 27, 28). Using the respective study’s cutoff for putative T2SS dependent effectors, we identified 16, 5, and 4 lipoproteins in the respective datasets. Upon compiling a list of putative T2SS dependent lipoproteins (**Table 1**), we cross-referenced these proteins against proteomic data sets of outer membrane vesicles (OMVs) produced by *A. baumannii* (29-32). We reasoned that the lipoproteins identified in the T2SS studies are most likely associated with OMVs given that their acyl groups facilitate membrane interaction. This has previously been demonstrated for the lipoprotein InvL (11). In agreement with this hypothesis, 15 of the 19 identified lipoproteins putatively secreted by the T2SS were also found in the proteomes of OMVs (**Table 1**). The function of most of these lipoproteins has yet to be determined; however, of the characterized proteins, FilF was previously identified as a cell surface antigen and promising vaccine target (33). The second characterized lipoprotein is InvL, an adhesin that depends on GspD for secretion and surface localization (11). Finally, RE180_13595 has been characterized as a glutamyl transferase and to be secreted in a GspD dependent manner, although the surface localization of this enzyme is still unclear (34).

**Table 1.**
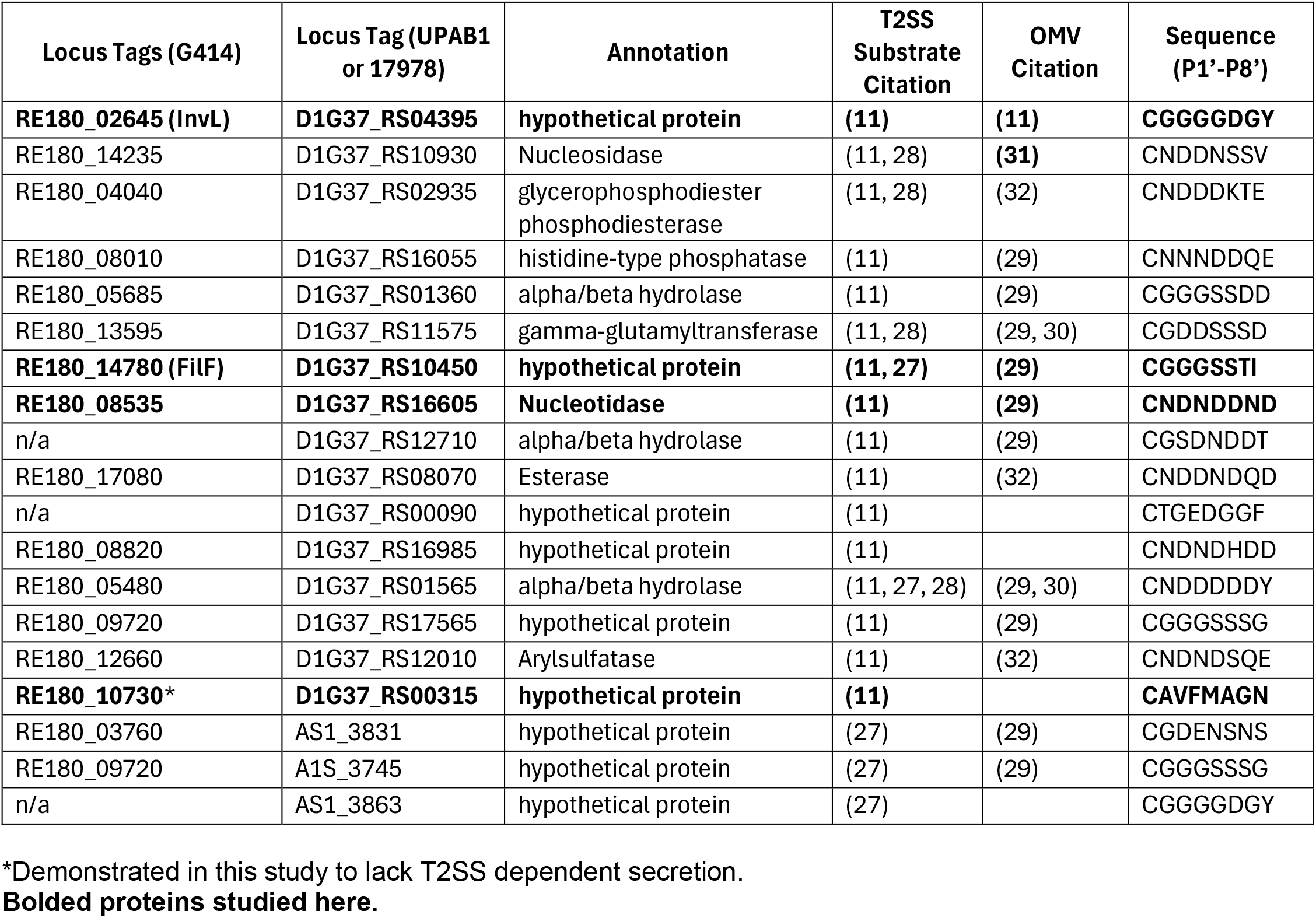
Verified or putative T2SS substrates with predicted lipobox.

We next analyzed the conservation of T2SS dependent lipoproteins genes across *A. baumannii* genomes, sorting the results by international clone (IC) groups (**Sup. Figure 2**). We excluded RE180_10730 from this analysis for reasons discussed below. Previously a similar analysis has been conducted for InvL that demonstrated high conservation across the IC groups except for IC1, IC6, and IC8, which we also observed here (11). As for the other lipoproteins, we found that the majority are conserved across all isolates regardless of IC group. For example, FilF is universal amongst all IC groups besides IC7 where it is still highly conserved. Three lipoproteins, RE180_03760 (IC2 and IC6), A1S_3863 (IC1, IC6 and IC8), and DIG37_RS00090 (IC7) are specific to certain IC groups. Overall, we found that all the strains examined except for one (CP169766.1), harbor ≥ 12 lipoproteins. Of the one outliner, this strain harbors the gene product for the putative gamma-glutamyl transferase (RE180_13595), which is universal across all strains examined.

### GspN is required for the secretion of InvL but not CpaA

Given the ubiquitous nature of the T2SS lipoprotein effectors and GspN in *A. baumannii* we next sought to experimentally validate a connection between the two. To address this, we first determined if these lipoproteins are encoded by a clinical strain that was recently isolated and sequenced (Gura et al, to be published elsewhere). We used SignalP prediction (35) to identify 150 putative lipoproteins encoded by G414, which we blasted against the putative T2SS dependent lipoproteins in **Table 1**. This revealed that 16 of the 19 identified lipoproteins are encoded by G414. We also identified genes for T2SS core components including *gspN* contained within an operon with *gspC* and *gspD*. For our studies, we chose to initially focus on the lipoprotein InvL, given its previous characterization and presence in G414. C-terminally histidine-tagged InvL has previously been shown to require GspD for surface localization (11). To test the potential involvement of GspN in the outer membrane translocation of InvL, we ectopically expressed InvL with a C-terminal His6 tag in wild-type (WT), Δ*gspD*, and Δ*gspN* mutant strains and examined secretion via western blotting of cell and culture supernatant fractions. As a control, we also analyzed the secretion of the well characterized and soluble His6-tagged T2SS substrate CpaA (6, 36-40). It should also be noted that we previously showed that the secretion of the T2SS effector LipA, a soluble lipase, is independent of GspN (2). Here, we found that while GspD is required for the secretion of both InvL and CpaA, only InvL depends on GspN (**Figure 2A**). We noted that there is slightly more CpaA in the cell pellet of the Δ*gspN* mutant as compared to the WT strain, but this is in stark contrast to the results observed with CpaA in the Δ*gspD* strain where all of CpaA is cell associated. Complementation of the Δ*gspN* mutant with genomically expressed GspN restored the secretion of InvL establishing an essential role for GspN in secretion of InvL but not the soluble protein CpaA (**Sup. Figure 3A**).

**Figure 2.**
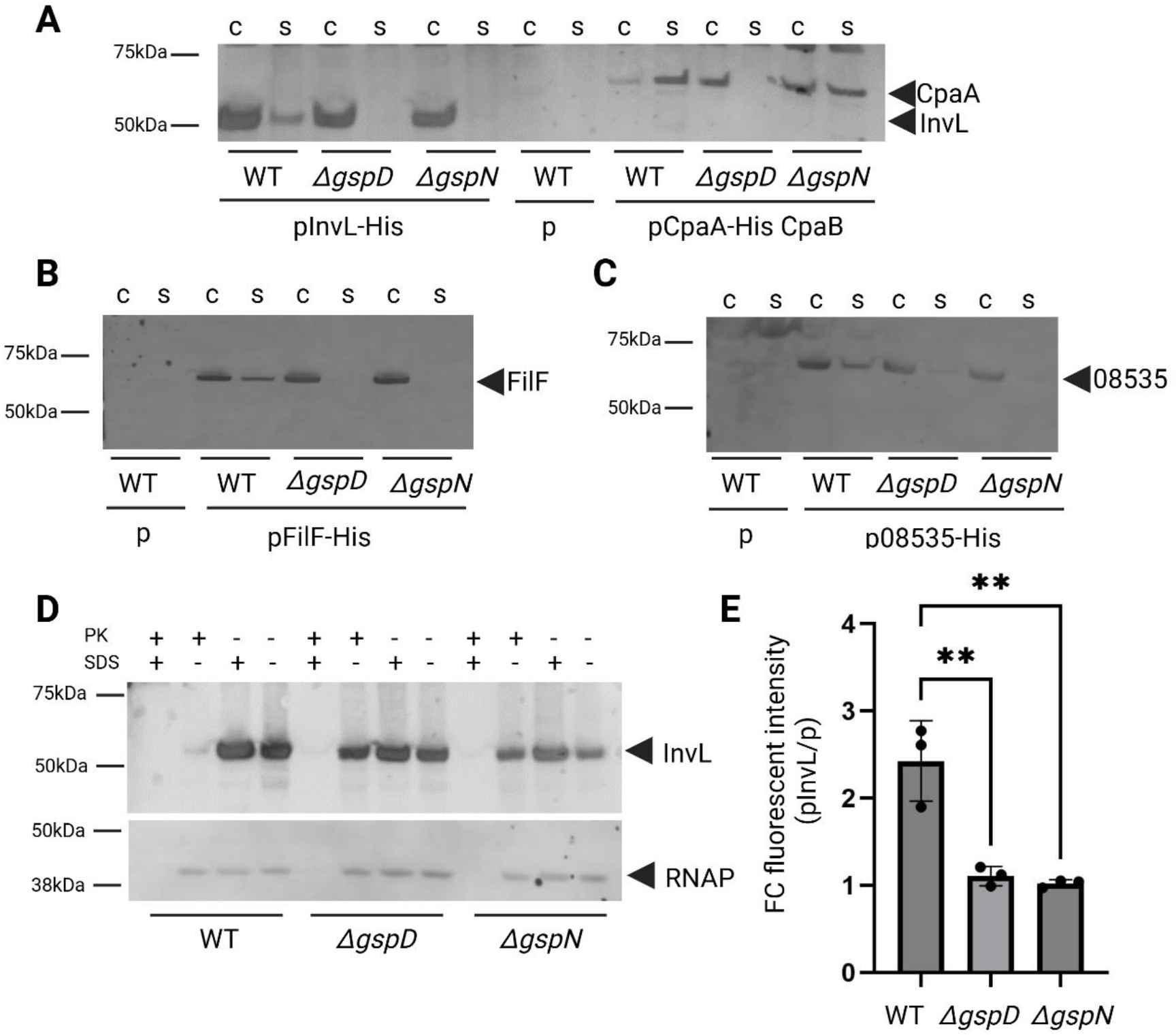
GspN is required for the secretion and surface localization of lipoprotein T2SS effectors. **A**. Overnight cultures from WT and indicated mutant strains containing empty vector (p) or plasmid expressing InvL or CpaA with C-terminal His6 tags were separated into cell (c) and supernatant (s) fractions, run on sodium dodecyl sulfate-polyacrylamide gel electrophoresis (SDS-PAGE), transferred to a nitrocellulose membrane, and blotted with anti-His6 antibody. Positions of molecular mass markers, InvL and CpaA are indicated. The plasmid encoding CpaA also codes for the chaperone CpaB to ensure proper folding and secretion of CpaA. WT with empty vector (p) was used as a negative control. **B**. Overnight cultures from WT and indicated mutant strains containing empty vector (p) or plasmid expressing FilF with a C-terminal His6 tag were assessed for FilF secretion as in **A**. Positions of molecular mass markers and FilF are shown. **C**. Overnight cultures from WT and indicated mutant strains with empty vector (p) or plasmid coding for 08535 with a C-terminal His6 tag were assessed for 08535 secretion as in **A**. Positions of molecular mass markers and protein 08535 are indicated. **D**. Cell pellets from overnight cultures from WT and indicated mutant strains expressing InvL with a C-terminal His6 tag were assessed for surface localization of InvL using proteinase K susceptibility with and without permeabilization of cells with 1% SDS. As a control, RNAP blots were stained with antibody against RNA polymerase α subunit. Positions of molecular mass markers, InvL and RNAP are indicated. **E**. Cell pellets from **D** were used to assess surface localization of InvL by incubating intact cells with His6 antibody followed by Alexa fluor 488 and monitoring the fluorescence intensity using excitation/emission wavelengths of 488/519 nm. Individual values represent relative fluorescence units of InvL expressing strain over strain containing empty vector. Data represent the average of three biological experiments with three technical replicates each +/-S.D. Tukey’s multiple comparison test was used with ** representing P < 0.005. Only significant differences are shown. In all cases, representative western blots are shown (n=2-3).

To determine if GspN is required for the secretion of additional T2SS dependent lipoproteins, we expressed both FilF and the putative nucleotidase RE180_08535 with a C-terminal His6 tag in WT, Δ*gspD*, and Δ*gspN* mutant strains and examined secretion via western blotting (**Figure 2B and C**). A similar banding pattern to InvL was observed for both proteins where the WT strain was shown to secrete FilF and RE180_08535, but the Δ*gspD* and Δ*gspN* mutant strains were not. Given the association with OMVs for InvL, FilF and RE180_08535 (11, 29), we also subjected the culture supernatant from WT cells expressing these proteins to filtration and ultracentrifugation to pellet crude OMVs. In all three cases, we observed that the majority of lipoprotein detected in the culture supernatant co-pelleted at high speeds with the lipid content found in the culture supernatant as assessed with the fluorescent dye FM 4-64 (**Sup. Figure 4**). This suggests that all three proteins are associated with WT OMVs. Together, these results indicate that GspN is required for the secretion of a subset of T2SS effectors in *A. baumannii* to the cell surface, which are then released to the extracellular space via OMVs.

### GspN is required for surface localization of InvL

We noted that in the case of InvL, FilF and RE180_08535, the majority of the protein is present in the cell pellet when ectopically expressed even in the WT strain. Previously this fraction of InvL has been shown to be surface localized (11). To determine if GspN is required to direct InvL to the cell surface, we subjected intact cells of WT, Δ*gspD*, and Δ*gspN* mutant strains expressing InvL to proteinase K accessibility. We observed that for the WT strain, InvL is largely surface localized given its sensitivity to proteinase K digestion in the absence of the permeabilizing reagent sodium dodecyl sulfate (SDS) (**Figure 2D**). In agreement with previously published findings, InvL is protected from proteinase K degradation in the Δ*gspD* mutant strain in the absence of SDS (11). A similar banding pattern to the Δ*gspD* mutant was observed for InvL expressed in the Δ*gspN* strain, indicating that GspN is required for the surface localization of InvL. To verify the role of GspN, we performed the same experiment with the *gspN* complemented Δ*gspN* strain, and again, observed near complete proteinase K susceptibility in the absence of SDS (**Sup. Figure 3B**). Using a quantitative approach, we also subjected intact cells to antibody labeling and measured the fluorescence intensity against His6-tagged InvL using anti-His6 antibody and Alexa fluor 488 conjugated secondary antibodies (**Figure 2E, Table 2**). We found that when InvL was expressed in WT cells, we observed significantly increased fluorescent intensity as compared to Δ*gspD* and Δ*gspN* mutant cells expressing InvL in agreement with our proteinase K sensitivity-based results.

**Table 2.** Relative fluorescence intensity of cells containing indicated plasmid over cells containing empty vector.

|  | **WT** | **$\Delta gspD$** | **$\Delta gspN$** |
| --- | --- | --- | --- |
| **pInvL** | 2.42 ± 0.37 | 1.10 ± 0.09 | 1.02 ± 0.03 |
| **pSP<sup>Blac</sup>-InvL** | 0.97 ± 0.07 | 0.94 ± 0.04 | 0.85 ± 0.10 |
| **p10730** | 1.03 ± 0.10 | 1.02 ± 0.08 | 0.89 ± 0.05 |
| **pSP<sup>Pal</sup>-InvL** | 1.04 ± 0.07 | 1.11 ± 0.07 | 1.04 ± 0.08 |
| **pSP<sup>LirL</sup>-InvL** | 0.92 ± 0.09 | 0.98 ± 0.05 | 0.93 ± 0.22 |
| **pSP<sup>08535</sup>-InvL** | 2.16 ± 0.11 | 0.78 ± 0.04 | 0.98 ± 0.08 |
| **pPal** | 1.06 ± 0.03 | - | - |
| **pLirL** | 0.97 ± 0.11 | - | - |
| **pSP<sup>InvL</sup>-Pal** | 1.03 ± 0.12 | - | - |

Mean ± standard deviation (n = 3).

To determine if GspN dependent outer membrane translocation and surface localization of InvL requires its lipobox, we generated a chimeric InvL species by replacing its own signal peptide with that of the normally periplasmic protein β-lactamase. First, we used SignalP 6.0 to predict the signal peptides and respective cleavage sites of the β-lactamase and InvL precursors (35). Then we substituted the signal peptide of InvL with the signal peptide from β-lactamase. This construct, SPβlac-InvL, was expressed in the WT cells and subjected to proteinase K treatment (**Sup. Figure 5A**). We found that this chimeric protein was protected from proteinase K digestion in the absence of permeabilizing reagent. We also quantitatively validated that SPβlac-InvL is not surface localized by performing surface labeling and quantitation using Alexa fluor 488 secondary antibodies (**Table 2**). To further confirm the intracellular location and loss of membrane interaction for InvL when expressed in the absence of its native lipobox, we subjected WT cells expressing SPβlac-InvL to subcellular fractionation (41). As a control for periplasmic content, we assessed each fraction for β-lactamase activity using a nitrocefin assay. SPβlac-InvL primarily associated with the β-lactamase activity-containing periplasmic fraction (**Sup. Figure 5B**). These findings support the previous conclusion (11) that InvL is indeed a lipoprotein and uses its lipobox for membrane association. We note that a small amount of SPβlac-InvL was released to the culture supernatant in both the WT and Δ*gspN* mutant strains, but not in the Δ*gspD* mutant (**Sup. Figure 5C**).

### A specific sorting motif is required but not sufficient for T2SS mediated secretion

Lipoproteins are often sorted to their final destinations in Gram-negative bacteria by a specific sequence C-terminal to the acylated cysteine residue of the lipobox but sorting rules vary across species (42). We reasoned that lipoproteins destined for T2SS-mediated surface localization might have a unique sequence motif that facilitates their trafficking to the cell surface. Previously, a cell surface signal was identified in the *Bacteroides* phylum that can heterologously target proteins to the cell surface (43), while another distinct signal was identified in proteins enriched in OMVs (44). Using the predicted lipobox sequences of the T2SS effectors expressed in G414 from **Table 1** we generated a sequence logo (45) starting with the universal cysteine residue at position P1’ relative to the signal peptidase II cleavage site and ending at P8’ (**Figure 3A**). This analysis revealed the enrichment of glycine, serine, aspartate, and asparagine residues, which we used as a cutoff to generate chimeric constructs in this study. Manual examination revealed that putative T2SS dependent lipoproteins are sorted into two groups where members contain either a glycine/serine rich motif (>4 amino acids) or a motif rich in asparagines/aspartates (>4 amino acids). Sequence logos generated from the sorted groups revealed a strong ‘GGG’ or ‘ND(N/D)’ motif at position P2’-P4’. We next generated a sequence logo for the remaining predicted 134 lipoproteins found encoded within G414’s genome at the same positions. This analysis did not reveal any clear sequence motif (**Figure 3B**), which may be due to competing sorting motifs between inner and outer membrane localized lipoproteins. To sort any potential distinct motif, we predicted the cellular localization of this group of lipoproteins using Psort (46) yielding 12 proteins predicted to localize to the inner membrane (**Table S1**), 17 predicted to localize to the outer membrane (**Table S2**), with the remaining predicted to localize to the extracellular space, periplasmic space, or had a low confidence prediction. Selecting the two membrane subsets to make specific sequence logos again did not result in a clearly recognizable motif at the same position highlighting the unique signal identified in T2SS dependent lipoproteins.

**Figure 3.**
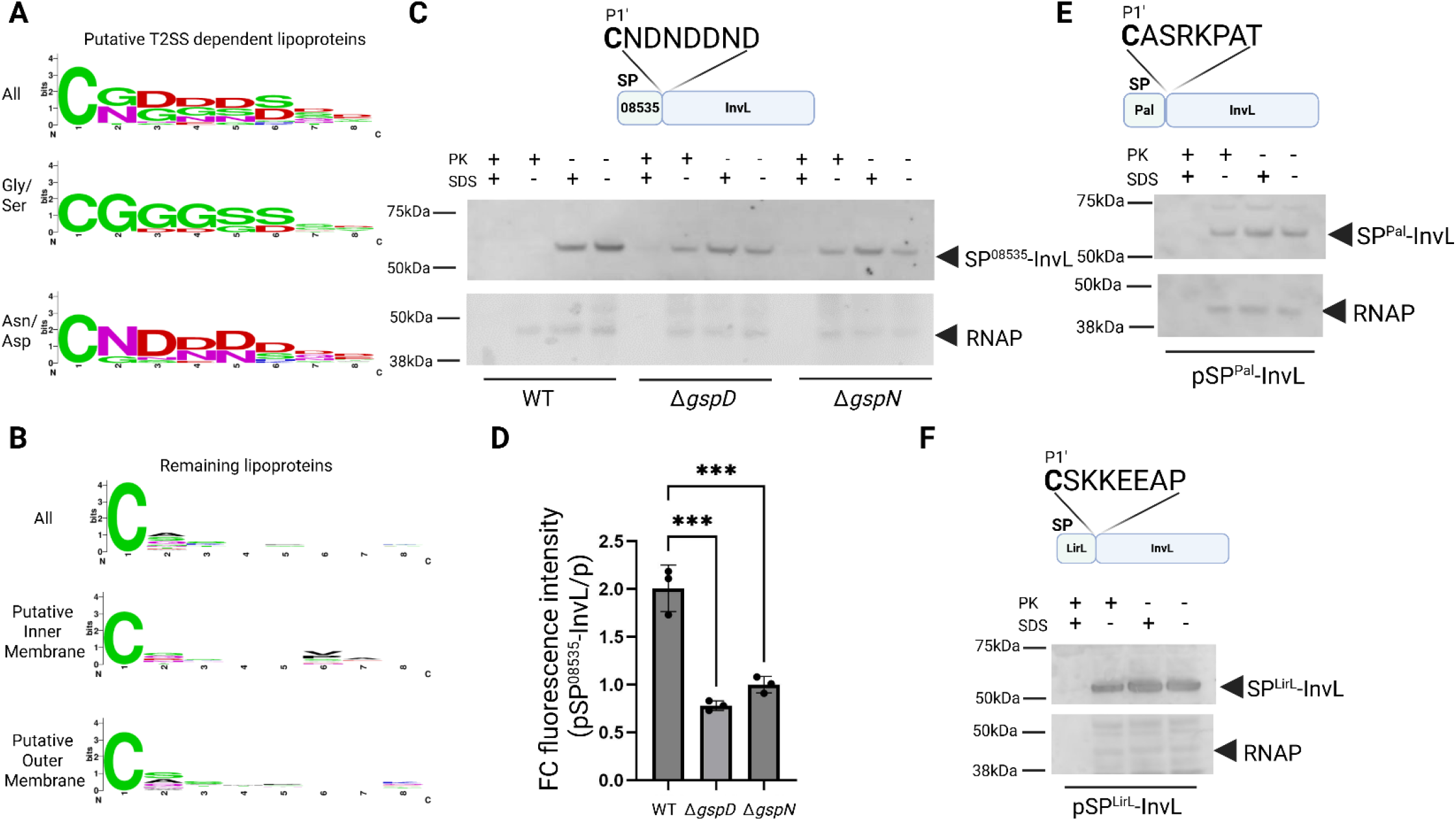
A specific sorting motif is present in lipoprotein T2SS effectors of *A. baumannii* and required for surface localization. **A**. Sequence logo generated with WebLogo showing residues P1’-P8’ of all putative T2SS dependent lipoproteins identified in **Table 1** with the exception of 10730. Sequences were further separated into glycine/serine rich and asparagine/aspartate rich, with resulting logos represented. **B**. A similar sequence logo was generated for the remaining lipoproteins predicted form the G414 genome. Subcellular localization was predicted using Psort to subsequently generate inner membrane and outer membrane sequence logos (list of proteins used found in **Table S1** and **S2** respectively). **C**. Schematic of signal peptide InvL chimera in which the signal peptide of InvL was replaced with that of 08535 to generate SP^08535^-InvL where P1’ – P8’ residues represented in relation to predicted signal peptidase II cleavage for 08535 (top). Cell pellets from overnight cultures were assessed for surface localization of SP^08535^-InvL using proteinase K susceptibility with and without permeabilization of cells with 1% SDS. As a control, blots were stripped and re-stained with antibody against RNA polymerase α subunit. Positions of molecular mass markers, SP^08535^-InvL and RNAP are indicated. Representative western blots (n=2-3). **D**. Cell pellets from **C** were used to assess surface localization of SP^08535^-InvL by labeling with His6 antibody followed by Alexa fluor 488 and monitoring the fluorescence intensity using excitation/emission wavelengths of 488/519 nm. Individual values represent fold change (FC) of relative fluorescence units (RFUs) of SP^08535^-InvL-expressing strain over empty vector containing strain where an average +/-S.D. of three biological experiments were used with three technical replicates. Tukey’s multiple comparison test used with *** representing P < 0.0005. Only significant differences are shown. **E**. Schematic of signal peptide InvL chimera in which the signal peptide of InvL was replaced with that of Pal to generate SP^Pal^-InvL where P1’ – P8’ residues represented in relation to predicted signal peptidase II cleavage site for Pal are indicated (top). Cell pellets from WT strain expressing the His6-tagged SP^Pal^-InvL were assessed for surface localization of SP^Pal^-InvL using proteinase K susceptibility with and without permeabilization of cells with 1% SDS. As a control, blots were stripped and re-stained with antibody against RNA polymerase α subunit. Positions of molecular mass markers, SP^Pal^-InvL and RNAP are shown. **F**. Schematic of signal peptide InvL chimera in which the signal peptide of InvL was replaced with that of LirL to generate SP^LirL^-InvL where P1’ – P8’ residues represented in relation to predicted signal peptidase II cleavage site for LirL are indicated (top). Cell pellets from WT strain expressing the His6-tagged SP^LirL^-InvL were assessed for surface localization of SP^LirL^-InvL using proteinase K susceptibility with and without permeabilization of cells with 1% SDS. As a control, blots were stripped and re-stained with antibody against RNA polymerase α subunit. Positions of molecular mass markers, SP^Pal^-InvL and RNAP are shown. In all cases, representative western blots are shown (n=2-3).

One putative T2SS dependent lipoprotein lacks an enrichment of glycine/serine or asparagine/aspartate residues, harboring a sequence motif of “CAVFMAGN” found in RE180_10730. Given the clear motif present in the remaining putative T2SS dependent lipoproteins, we chose to follow up on RE180_10730. To assess whether RE180_10730 is indeed secreted in a T2SS dependent manner, we expressed RE180_10730 with a C-terminal His6 tag in WT, Δ*gspD*, and Δ*gspN* mutant strains. We did not observe significant secretion of this protein as determined with western blotting in any of the strains tested despite detecting the protein in the cell fraction (**Sup. Figure 6A)**. We utilized proteinase K susceptibility to rule out that the cellular RE180_10730 is exposed on the cell surface (**Sup. Figure 6B**) and performed additional validation using surface labeling (**Table 2**). Furthermore, bioinformatic analysis indicated that RE180_10730 is likely an outer membrane protein that may contribute to the assembly of outer membrane proteins as it is homologous to SmpA, which forms part of the YaeT (BamA) complex in *E. coli* (47). We therefore speculate that RE180_10730 is a false positive identified in previous secretome studies. Secretome data could easily include proteins present in the inner leaflet of the outer membrane as this group of proteins can also be released though OMVs, although we did not see RE180_10730 in any of the published OMV proteomic data. It is also possible that the expression of RE180_10730 is for unknown reasons reduced in T2SS mutants of *A. baumannii*. We note that RE180_10730 was at the lower end of the cutoff threshold in its respective study (11).

After identifying a putative glycine/serine or asparagine/aspartate signal in T2SS dependent lipoproteins, we tested whether a signal peptide with an adjacent ND(N/D) motif from another GspN-dependent lipoprotein can support the surface localization of InvL which has a GGG motif by generating an InvL chimera consisting of the signal peptide, conserved cysteine, and seven adjacent residues from 08535. Like WT InvL, SP08535-InvL was secreted in the WT strain, but not Δ*gspD* and Δ*gspN* mutants (**Sup. Figure 7A**). We also verified the surface accessibility of this construct in WT and mutant cells by assessing resistance to proteinase K treatment. We observed that SP08535-InvL was sensitive to proteinase K treatment in the WT strain regardless of SDS-treatment, while this construct expressed in Δ*gspD* and Δ*gspN* mutants was protected in the absence of SDS (**Figure 3C**). In agreement, WT but not mutant cells expressing SP08535-InvL were efficiently labeled using His6-tag antibody and Alexa fluor 488 secondary antibodies (**Figure 3D, Table 2**). Next, we determined if the lipobox from an intracellular lipoprotein would be tolerable for secretion of InvL. Previously the lipoprotein Pal was identified as an outer membrane protein localized to the inner leaflet (48). To verify that Pal is not a secreted protein, we first expressed Pal with a C-terminal His6 tag in WT, Δ*gspD*, and Δ*gspN* mutant strains, separated cell and supernatant fractions and probed for Pal via western blotting. We did not observe significant secretion of Pal in any strain despite detection of a cell associated fraction (**Sup. Figure 8A**). We also verified that Pal is not exposed on the cell surface of intact cells as it was resistant to proteinase K treatment and did not exhibit an increase in fluorescence intensity upon surface labeling (**Sup. Figure 8B, Table 2**). After verifying that Pal is indeed not secreted via the T2SS, we generated a chimeric InvL protein, SPPal-InvL, containing the signal peptide, conserved cysteine, and seven adjacent residues from Pal. SPPal-InvL was found primarily associated with the cells and was protected from proteolysis by proteinase K in the absence of permeabilizing reagent (**Figure 3E, Sup. Figure 7B, Table 2**). In a similar approach, we generated a second chimera using the signal peptide, conserved cysteine, and seven adjacent residues from LirL, which is a lipoprotein previously found to localize to the inner membrane (49). After validating that LirL was not secreted or surface localized (**Sup. Figure 9, Table 2**), we probed the surface localization and secretion of SPLirL-InvL using proteinase K accessibility assay (**Figure 3F, Sup. Figure 7C, Table 2**). We found that SPLirL-InvL was primarily associated with cells and was protected from proteolysis in the absence of permeabilizing reagent. Taken together, we conclude that native sorting motifs within the T2SS lipoproteins of *A. baumannii* are permissible for surface localization of T2SS dependent lipoproteins but not a sequence from the intracellular proteins such as Pal or LirL.

After verifying that InvL requires a specific motif for secretion we sought to determine if the lipobox from InvL is sufficient to guide a T2SS independent lipoprotein to the extracellular space. To this end, we generated a chimeric protein consisting of the lipobox, conserved cysteine, and seven adjacent residues from InvL attached to Pal (SPInvL-Pal). When we expressed this protein in WT, Δ*gspD*, and Δ*gspN* strains we primarily observed it in the cell fraction (**Sup. Figure 8C**). SPInvL-Pal in the WT cell fraction was protected from proteinase K in the absence of permeabilizing reagent and did not exhibit an increase in fluorescent labeling with His6-antibody and Alexa fluor 488 secondary antibodies compared to cells with empty vector, indicating that SPInvL-Pal is not surface localized (**Sup. Figure 8D, Table 2**). We therefore conclude that in addition to a specific sorting motif adjacent to the lipobox, lipoproteins destined for T2SS mediated surface localization also require a distinct secretion signal and that the specific sorting motif from InvL is required but not sufficient for surface localization.

### A sorting motif is also present in T2SS dependent lipoproteins in *B. pseudomallei and S. oneidensis*

Given the presence of GspN homologs in *B. pseudomallei* and *S. oneidensis*, we asked if these T2SS dependent lipoproteins also harbor a sorting motif similar to that of InvL and 08535. Several studies have characterized the functionality of the T2SS from *B. pseudomallei*, but a lipoprotein effector has yet to be directly characterized (50, 51). There are also some reports that have characterized the GspD dependent surface lipoproteins from *S. oneidensis* and showed that they can be used in bioenergy and bioremediation applications (52, 53). Taking advantage of published proteomic datasets, we analyzed the signal peptides of the T2SS dependent lipoproteins from each of these organisms using SignalP (35)(**Tables S3** and **S4**). Sequence logos (45) showed a similar sorting motif beginning at the P1’ position relative to signal peptidase cleavage site as compared to the sequence logo generated from all of the T2SS dependent lipoproteins from *A. baumannii* (**Sup. Figure 10A and B**) for both species, albeit the conservation was certainly stronger for *B. pseudomallei*. Manual examination of the individual sequences revealed that most putative T2SS dependent lipoproteins in these organisms harbor a hybrid motif mostly consisting of glycine and aspartate. This is in contrast to the bifurcated motif found in *A. baumannii*. Given the GspD dependence of these lipoproteins and a conserved sorting motif, we reason that these lipoproteins might also depend on GspN for surface localization.

### Pullulanase depends on PulN for surface localization in *K. pneumoniae*

Considering our results in *A. baumannii*, we sought to directly establish if T2SS dependent lipoproteins expressed by other bacteria similarly rely on GspN homologs for secretion. We focused on the lipoprotein PulA that is secreted by the T2SS in *K. pneumoniae* to the cell surface and is responsible for degradation of the polysaccharide pullulan (54). We assessed the ability of intact cells from WT and two transposon mutant strains with disruption of either *pulA* or *pulN (gspN* homolog) (55) of *K. pneumoniae* KPPR1 to degrade pullulan. Upon incubation of cells with pullulan, we found that the WT and *pulN*::Tn mutant strain complemented with ectopic *pulN* were able to degrade pullulanase as assessed by measuring the rate of reduced sugar generation (**Figure 4**). In contrast, the *pulA*::Tn and *pulN*::Tn mutant strains harboring empty vector had significantly lower pullulanase activity. When the same cells were evaluated for pullulanase activity after sonication, we observed significant pullulanase activity in the WT and *pulN*::Tn mutant strain regardless of plasmid complementation. This demonstrates that PulA is produced but not localized to the cell surface in cells with disruption of *pulN*. Consistent with the role of PulA in pullulan degradation, the *pulA*::Tn mutant strain did not have pullulanase activity even after sonication. These data indicate that PulN facilitates the surface localization of PulA. We note that the sorting motif of PulA differs from that in *A. baumannii* but is still enriched in serine residues. Collectively, a similar mechanism is suggested to govern the T2SS dependent secretion of lipoproteins to the cell surface across multiple organisms.

**Figure 4.**
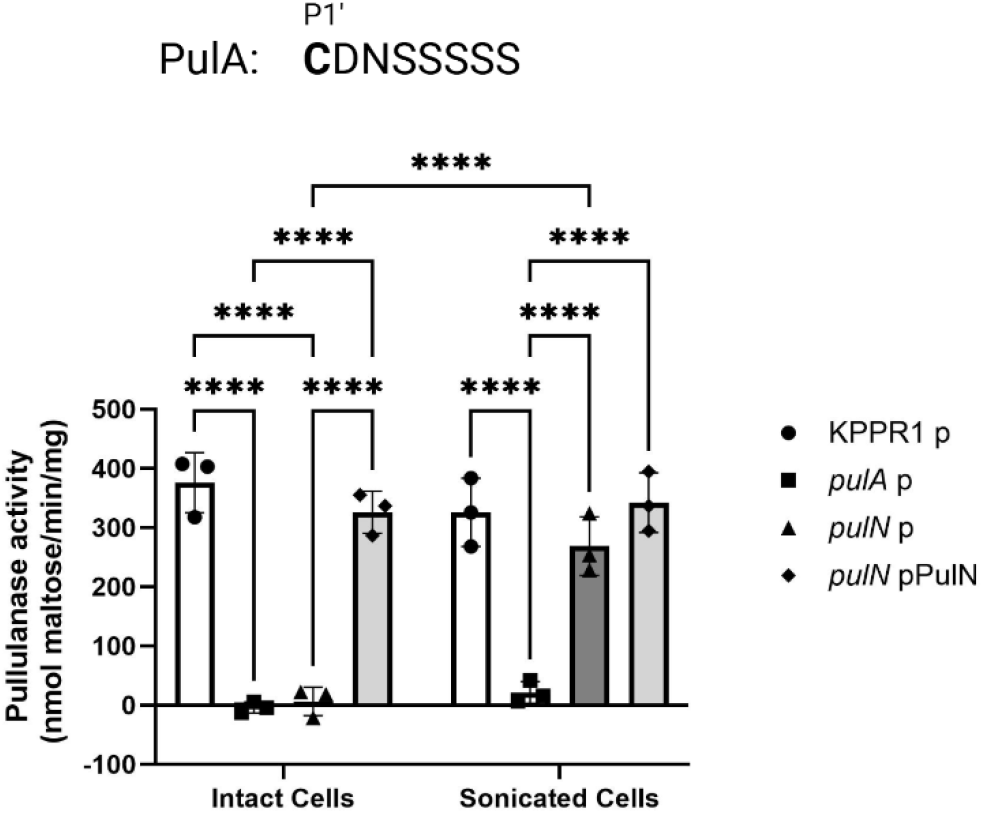
PulN is required for secretion of pullulanase. The sequence starting at P1’ relative to signal peptidase II cleavage site of the *pulD* (*gspD* homolog) dependent lipoprotein PulA from *K. pneumoniae* (top). Intact and sonicated cells from logarithmic phase cultures of *K. pneumoniae* strain KPPR and mutants with transposon insertions containing empty vector (p) or ectopically expressing PulN (GspN homolog) were isolated and assessed for pullulanase activity by monitoring the reduced sugar content produced over time against a maltose standard curve as determined with 3,5-dinitrosalicylic acid. Experiments were performed in biological triplicates with technical triplicates. Data are presented as mean +/-S.D. \*\*\*\**P* < 0.0001 by one-way ANOVA analysis with Šídák’s multiple comparisons test. Only significant differences are shown.

## Discussion

In this report, we have demonstrated, for the first time, GspN dependent secretion of T2SS effectors containing a lipobox with a specific sorting motif in *A. baumannii*. Only some gram-negative bacteria that encode a T2SS system express GspN, but the gene product is highly conserved in these species supporting a role in the specialized function in the transport of lipoproteins. Structural prediction reveals a relationship between GspN and members of the AsmA-like protein family known for their roles in phospholipid transfer between the inner and outer membranes in gram-negative bacteria. The repeating β-taco folds (also called β-groove motifs) of these proteins are thought to shield phospholipids from unfavorable interactions in the aqueous periplasmic compartment (18). In the context of the T2SS, we believe that the hydrophobic β-groove motif of GspN may shield the acyl chains of lipoproteins from unfavorable interactions, and we have demonstrated a genetic link in support of this hypothesis. Given its likely inner membrane localization, we propose that GspN initially recognizes lipoproteins destined for the cell surface after their posttranslational modification. While bacterial AsmA-like proteins have been indicated in maintaining phospholipid homeostasis between the inner and outer membrane (18), there has yet to be a report showing a role in the transport of additional molecules such as proteins. The six identified AsmA-like proteins found in *E. coli* greatly vary in predicted length from 132-268 Å. On the larger end, YhdP is believed to span the entirety of the periplasm, while smaller AsmA-like proteins such as TamB are known to interact with outer membrane proteins such as TamA to bridge the periplasmic gap (18). It seems likely that the latter case is also true for GspN based on its predicted globular length of only 60 Å. Additional T2SS components and perhaps other thus far unidentified proteins may aid in transporting lipoproteins possible via a hand-off mechanism from GspN. Alternatively, a soluble chaperone similar in function to LolA could be involved. Other proteins of the T2SS apparatus facilitate specialized function such as GspC that, depending of the species, contains a PDZ domain or a coiled-coil domain that is implicated in extracellular secretion of specific substrates (56-58). Here, we show that GspN has a specialized function in the transport of lipoproteins.

Canonically, the residues adjacent to the acylated cysteine of lipoproteins dictate their subcellular location. For example, in *E. coli* an aspartate residue is often seen as an inner membrane retention signal (59). However, inner membrane retention signals differ among gram-negative species. (60). In *Pseudomonas*, a ‘GGG’ or ‘GDD’ motif serves as avoidance signals for the Localization of lipoprotein (Lol) pathway, a transport pathway responsible for inserting lipoproteins in the inner leaflet of the outer membrane, resulting in inner membrane retention (60). The presence of amino acids serine or lysine also influence sorting in Pseudomonas (61). More recently, a unique sorting motif was found in *Bacteroides* lipoproteins that is sufficient for surface localization (43). In this study, bioinformatic analysis revealed the presence of a specific, bifurcated sorting motif inT2SS dependent lipoproteins of *A. baumannii* that differs from that in intracellular lipoproteins. This sorting motif is required, but not sufficient for outer membrane transport and surface localization of InvL. When the T2SS dependent lipoprotein InvL was expressed with the corresponding sequence from the normally intracellular Pal or LirL, the resulting constructs were no longer surface localized. Excitingly, the T2SS sorting motif is also present in putative T2SS dependent lipoproteins in other species, and we were able to use this sequence to correctly predict lipoproteins that are not secreted by the T2SS in *A. baumannii*. We speculate that the sorting motif in T2SS dependent lipoproteins allows both for avoidance of the Lol pathway and efficient recognition by GspN. Previously, PulA from *K. pneumoniae* was found to localize to the inner leaflet of the outer membrane when the aspartate at P2’ was substituted with other residues promoting recognition by the Lol pathway (62). We speculate that on top of a specific sorting motif, additional secondary structural information is required for outer membrane translocation. This is supported by the small, but detectable secretion of SPβlac-InvL.

While not all bacteria containing a T2SS express a GspN homolog, both *K. pneumoniae* and *V. cholerae* express GspN. Seminal work from the Puglsey group identified the genes required for secretion of the lipoprotein PulA from *K. pneumoniae* (3, 13, 14, 62, 63). While initially PulN was believed to play a role in the biogenesis of PulA (3), it was subsequently shown to be dispensable for PulA secretion in a later study (14). A key distinction between these two studies by the Puglsey group was the identity of the PulA species used in each study. In the first study, native PulA, was analyzed which contains a lipobox allowing for surface localization. Subsequently, a non-acylated version of PulA was used to identify essential genes for PulA secretion to the extracellular space. We would expect that the non-acylated variant of PulA to be secreted to the culture medium independent of *pulN* expression. PulA is thus far the only reported protein to rely on the T2SS in *K. pneumoniae*. While many T2SS substrates have been identified for *V. cholerae*, including the disease-causing cholera toxin, a bon a fide lipoprotein T2SS effector has yet to be confirmed. Perhaps additional T2SS dependent lipoproteins are produced by both these species but primarily localize to the cell surface explaining how they might have been missed in traditional secretome analyses. From the work presented here, we believe a repertoire of surface associated lipoproteins that previously showed GspD dependence also depend on GspN for secretion in notable pathogen *B. pseudomallei* and likely rely on a similar transport mechanism. In addition, *S. oneidensis*, which is being used for biotechnology applications, relies on a group of lipoproteins for electron transfer reactions that have previously been shown to depend on GspD for surface localization and likely also rely on GspN. In *A. baumannii*, T2SS dependent lipoproteins makes up a significant portion of the total number of lipoproteins (13%) and a large portion of the proteins secreted by the T2SS. Regarding UPAB1, a urinary tract isolate studied previously, over 50% of the putative T2SS dependent proteins are lipoproteins (11). Recently, the lipoprotein studied here, InvL, was shown to be important for establishing a chronic respiratory infection (64). Of the few additionally characterized lipoproteins identified in this study to be GspN dependent, the gamma-glutamyl transferase (RE180_13595) has been recognized as a virulence factor (34). These findings demonstrate the importance of further study of this newly identified mechanism of the T2SS.

## Materials and Methods

### Bacteria strains and plasmids

The *Acinetobacter baumannii* strain G414 (CP133499) is a modern clinical isolate from the University of Michigan Hospital. Isogenic mutants Δ*gspD* and Δ*gspN* were generated as previously described (2). The *gspN* complemented strains were generated by overlap extension PCR using genomic G414 DNA and plasmid DNA form pGspN (2) as template with the indicated primers before ligating into pCVD442. The *Klebsiella pneumoniae* strain used was KPRR1 and indicated mutant derivatives were generated previously (55). Additional plasmids and primers used in this study are listed in **Table S5**. All polymerase chain reactions (PCR), cloning, and restriction enzyme digestions were done with either SuperFi Platinum polymerase or PfuTurbo, T4 DNA ligase, and restriction enzymes from New England Biolabs and primers that were synthesized at IDT Technologies. Genes of interest were routinely cloned with the indicated primer using genomic G414 DNA as template before cloning into the Agilent Blunt End Vector Kit per manufacturer’s recommendations and mobilization into pMMB67EH for expression. pMMB67EH constructs were transformed into *E. coli* MC1061 and pCVD442 constructs into SY327λpir. Triparental conjugation was performed with a helper strain carrying pRK2013 to transfer plasmids into G414. Plasmids were mobilized into KPRR1 via electroporation. Indicated primers were used to generate the signal peptide chimeric proteins along with Fwd and Rev pMMB67EH primers where template DNA constituted plasmid DNA coding for the two proteins of interest before ligation back into pMMB67EH.

### Growth Conditions

Strains were grown on Luria-Bertani (LB) agar/broth (Fisher) with 100 µg/mL of carbenicillin (Sigma) when plasmids were present unless otherwise indicated. IPTG was added at a concentration of 50 µM to induce the expression of InvL and chimeras, CpaA, and Pal. IPTG was omitted from the growth media of strains expressing LirL, FilF, 08535, and 10730. For growth of *K. pneumoniae* strains, overnight cultures were started in LB broth followed by an outgrowth in M9 salts supplemented with 0.4% casamino acids and 0.4% maltose until an OD of 0.9 was reached.

### Sodium dodecyl sulfate-polyacrylamide gel electrophoresis and immunoblotting

Overnight cultures were normalized to an OD 600 of 2.5 before separating cells from supernatant fractions by centrifugation at 10,000 × g for 10 minutes. Fractions were analyzed by sodium dodecyl sulfate-polyacrylamide gel electrophoresis and immunoblotting as described previously. Antibodies against 6-His and the α subunit of RNAP were incubated with membranes for 2 h (1:5,000). Following washing, horseradish peroxidase-conjugated goat anti-rabbit immunoglobulin G or goat anti-mouse immunoglobulin G (Bio-Rad, Thermo Fisher) used at 1:20,000 were incubated with the membrane for 1 h. Membranes were developed using ECL 2 Western blotting reagent (Thermo Fisher) and visualized using a Typhoon V variable mode imager system and analyzed with ImageJ imager software.

For proteinase K accessibility assays, stationary phase cultures were resuspended in LB to an OD at 600 nm of 2.5. Suspensions were treated with 200 μg/mL proteinase K (11) or an equal volume of LB in the presence or absence of 1% SDS at 37°C for 30 min. Phenylmethylsulfonyl fluoride was added at 1 mM to stop the digestion, followed by SDS-PAGE and western blotting analysis as described above.

### Cell surface detection of His6-tagged proteins

Cells were washed, blocked with 2% BSA, and incubated with 1:1,000 of antibody against His6. Following incubation with 1:1,000 of Alexa Fluor 488 F(ab′)2 goat anti-rabbit immunoglobulin G (Invitrogen) and washing, fluorescence was measured (Ex 488 nm/Em 525 nm). The results were normalized to the fluorescence intensity of the same strain carrying an empty vector as described (58).

### Crude outer membrane vesicle preparation

Cells were grown overnight before culture supernatants were isolated by centrifugation at 10,000 × *g* for 10 minutes. Subsequent sterile filtration and centrifugation at 200,000 × *g* for 3 h pelleted the insoluble fraction. Separately, the cleared supernatant and the pellets containing crude outer membrane vesicles resuspended in LB were assayed via western blotting using 6-his antibody as described above. As a control, the fluorescent dye FM 4-64 was incubated with each fraction for 10 minutes in the dark, before monitoring the fluorescent intensity with excitation/emission wavelengths of 558/734 nm as described (41). Background intensity from media alone was first subtracted before normalizing to the intensity in total supernatant fractions.

### Subcellular fractionation

Log phase WT cells were harvested before resuspending in buffer containing 50 mM Tris pH 7.5, 100mM Sucrose, and 1 mM EDTA. Lysozyme was added to a final concentration of 1 mg/mL before incubation 5 minutes at room temperature. Samples were centrifuged at 10,000 × g for 15 minutes, and the resulting supernatant was collected as the periplasmic content. The remaining pellet was resuspended in the same buffer with the addition of 1 mM PMSF and 10 µg/mL DNAse. Cells were sonicated with a probe type sonicator. After removing the cell debris, the cytoplasmic and membrane content was separated by ultracentrifugation at 100,000 g for 1 hour. Subsequent supernatant fraction was saved as cytoplasmic content, while the pellet was resuspended in the same buffer and taken as membrane content. As a control, β-lactamase activity was assessed with nitrocefin as described (65). Activity was determined by monitoring the change in absorbance at 500 nm and subtracting by the change in absorption of buffer alone. Cytoplasmic, periplasmic, and membrane activities were then added up and the individual activities were reported as a percentage of the total from the three.

### Assessment of pullulanase activity

Total and surface pullulanase activity was assessed as previously described (66). Briefly, cultures were pelleted, washed three times in phosphate buffered saline (PBS) before intact cells were resuspended in PBS and incubated with 10 mg/mL pullulan for 4 hours. Aliquots were withdrawn at intervals and assessed for reduced sugars using dinitrosalicylic acid against a maltose standard curve. Pullulan activity is expressed as nanomole maltose produced per minute per milligram of total protein as assessed by Bicinchoninic Acid assay. To compare surface versus trapped pullulan activity, cells were also sonicated prior to pullulan incubation. As a control, pullulan was incubated with PBS alone. Assays were performed in ≥3 biological replicas in technical triplicates. One-way ANOVA with a Šídák’s correction for multiple comparisons was used to compare all values. Results yielding a *P* value of <0.05 were considered statistically significant; values >0.05 were not shown. All calculations were done using GraphPad Prism version 10.0.0 for Windows, GraphPad Software, Boston, MA, USA; https://www.graphpad.com).

### Structural analysis, sequence alignment, and bioinformatic analysis

The primary amino acid sequence of GspN from G414 was used to predict the secondary structure with AlphaFold3 (67). The predicted structure of PulN, EpsN, and AsmA was downloaded from the AlphaFold database. Structures were visualized with ChimeraX. For the sequence logos of G414 lipoproteins, the 8 amino acids C-terminal to signal peptidase II cleavage site were used to generate a logo using the Berkley WebLogo online tool. For prediction of signal peptides, SignalP 6 (35) was used. For gene conservation, complete genomes were extracted from NCBI to form a custom database and blasted against gspN, *pulN*, and *epsN* for *A. baumannii, K. pneumoniae*, and *V. cholerae* respectively using command line blast. Lipoproteins were similarly blasted against complete *A. baumannii* genomes. Results were subsequently divided into IC groups using a custom script.

## Acknowledgments

We would like to thank Drs. Catilyn Holmes and Michael Bachman for their kind gift of WT and mutant strains of *K. pneumoniae* strain KPRR1. This work was, in part, supported by R01AI137085 (to M.S.) and F32AI178852 (to C.R.).

## References

1. K. V. Korotkov, M. Sandkvist, Architecture, Function, and Substrates of the Type II Secretion System. EcoSal Plus 8 (2019).

2. T. L. Johnson, U. Waack, S. Smith, H. Mobley, M. Sandkvist, Acinetobacter baumannii Is Dependent on the Type II Secretion System and Its Substrate LipA for Lipid Utilization and In Vivo Fitness. J Bacteriol 198, 711–719 (2015).

3. I. Reyss, A. P. Pugsley, Five additional genes in the pulC-O operon of the gram-negative bacterium Klebsiella oxytoca UNF5023 which are required for pullulanase secretion. Mol Gen Genet 222, 176–184 (1990).

4. A. E. Sikora, R. A. Zielke, D. A. Lawrence, P. C. Andrews, M. Sandkvist, Proteomic analysis of the Vibrio cholerae type II secretome reveals new proteins, including three related serine proteases. J Biol Chem 286, 16555–16566 (2011).

5. M. Sandkvist, V. Morales, M. Bagdasarian, A protein required for secretion of cholera toxin through the outer membrane of Vibrio cholerae. Gene 123, 81–86 (1993).

6. U. Waack et al., CpaA Is a Glycan-Specific Adamalysin-like Protease Secreted by Acinetobacter baumannii That Inactivates Coagulation Factor XII. mBio 9 (2018).

7. C. S. Roberts, A. B. Shannon, K. V. Korotkov, M. Sandkvist, Differential processing of VesB by two rhomboid proteases in Vibrio cholerae. mBio 15, e0127024 (2024).

8. C. d’Enfert, C. Chapon, A. P. Pugsley, Export and secretion of the lipoprotein pullulanase by Klebsiella pneumoniae. Mol Microbiol 1, 107–116 (1987).

9. Y. Ferrandez, G. Condemine, Novel mechanism of outer membrane targeting of proteins in Gram-negative bacteria. Mol Microbiol 69, 1349–1357 (2008).

10. A. L. Horstman, M. J. Kuehn, Bacterial surface association of heat-labile enterotoxin through lipopolysaccharide after secretion via the general secretory pathway. J Biol Chem 277, 32538–32545 (2002).

11. C. D. Jackson-Litteken et al., InvL, an Invasin-Like Adhesin, Is a Type II Secretion System Substrate Required for Acinetobacter baumannii Uropathogenesis. mBio 13, e0025822 (2022).

12. S. Gadwal, T. L. Johnson, H. Remmer, M. Sandkvist, C-terminal processing of GlyGly-CTERM containing proteins by rhombosortase in Vibrio cholerae. PLoS Pathog 14, e1007341 (2018).

13. A. P. Pugsley, I. Reyss, Five genes at the 3’ end of the Klebsiella pneumoniae pulC operon are required for pullulanase secretion. Mol Microbiol 4, 365–379 (1990).

14. O. M. Possot, G. Vignon, N. Bomchil, F. Ebel, A. P. Pugsley, Multiple interactions between pullulanase secreton components involved in stabilization and cytoplasmic membrane association of PulE. J Bacteriol 182, 2142–2152 (2000).

15. A. A. Chernyatina, H. H. Low, Core architecture of a bacterial type II secretion system. Nat Commun 10, 5437 (2019).

16. S. Okuda, H. Tokuda, Lipoprotein sorting in bacteria. Annu Rev Microbiol 65, 239–259 (2011).

17. S. Kumar, N. Ruiz, Bacterial AsmA-Like Proteins: Bridging the Gap in Intermembrane Phospholipid Transport. Contact (Thousand Oaks) 6, 25152564231185931 (2023).

18. S. Kumar, R. M. Davis, N. Ruiz, YdbH and YnbE form an intermembrane bridge to maintain lipid homeostasis in the outer membrane of Escherichia coli. Proc Natl Acad Sci U S A 121, e2321512121 (2024).

19. M. V. Douglass, A. B. McLean, M. S. Trent, Absence of YhdP, TamB, and YdbH leads to defects in glycerophospholipid transport and cell morphology in Gram-negative bacteria. PLoS Genet 18, e1010096 (2022).

20. S. S. Abby et al., Identification of protein secretion systems in bacterial genomes. Sci Rep 6, 23080 (2016).

21. M. S. Decanio, R. Landick, R. J. Haft, The non-pathogenic Escherichia coli strain W secretes SslE via the virulence-associated type II secretion system beta. BMC Microbiol 13, 130 (2013).

22. M. Chami et al., Structural insights into the secretin PulD and its trypsin-resistant core. J Biol Chem 280, 37732–37741 (2005).

23. J. Arts et al., Interaction domains in the Pseudomonas aeruginosa type II secretory apparatus component XcpS (GspF). Microbiology (Reading) 153, 1582–1592 (2007).

24. S. R. Lybarger, T. L. Johnson, M. D. Gray, A. E. Sikora, M. Sandkvist, Docking and assembly of the type II secretion complex of Vibrio cholerae. J Bacteriol 191, 3149–3161 (2009).

25. H. M. Lee et al., Association of the cytoplasmic membrane protein XpsN with the outer membrane protein XpsD in the type II protein secretion apparatus of Xanthomonas campestris pv. campestris. J Bacteriol 182, 1549–1557 (2000).

26. R. C. White et al., Type II Secretion-Dependent Aminopeptidase LapA and Acyltransferase PlaC Are Redundant for Nutrient Acquisition during Legionella pneumophila Intracellular Infection of Amoebas. mBio 9 (2018).

27. N. M. Elhosseiny, O. M. El-Tayeb, A. S. Yassin, S. Lory, A. S. Attia, The secretome of Acinetobacter baumannii ATCC 17978 type II secretion system reveals a novel plasmid encoded phospholipase that could be implicated in lung colonization. Int J Med Microbiol 306, 633–641 (2016).

28. N. M. Elhosseiny, N. B. Elhezawy, A. S. Attia, Comparative proteomics analyses of Acinetobacter baumannii strains ATCC 17978 and AB5075 reveal the differential role of type II secretion system secretomes in lung colonization and ciprofloxacin resistance. Microb Pathog 128, 20–27 (2019).

29. Z. T. Li et al., Outer membrane vesicles isolated from two clinical Acinetobacter baumannii strains exhibit different toxicity and proteome characteristics. Microb Pathog 81, 46–52 (2015).

30. S. O. Kwon, Y. S. Gho, J. C. Lee, S. I. Kim, Proteome analysis of outer membrane vesicles from a clinical Acinetobacter baumannii isolate. FEMS Microbiol Lett 297, 150–156 (2009).

31. J. S. Jin et al., Acinetobacter baumannii secretes cytotoxic outer membrane protein A via outer membrane vesicles. PLoS One 6, e17027 (2011).

32. G. Dhurve, A. K. Madikonda, M. V. Jagannadham, D. Siddavattam, Outer Membrane Vesicles of Acinetobacter baumannii DS002 Are Selectively Enriched with TonB-Dependent Transporters and Play a Key Role in Iron Acquisition. Microbiol Spectr 10, e0029322 (2022).

33. R. Singh, N. Garg, G. Shukla, N. Capalash, P. Sharma, Immunoprotective Efficacy of Acinetobacter baumannii Outer Membrane Protein, FilF, Predicted In silico as a Potential Vaccine Candidate. Front Microbiol 7, 158 (2016).

34. N. M. Elhosseiny et al., gamma-Glutamyltransferase as a Novel Virulence Factor of Acinetobacter baumannii Inducing Alveolar Wall Destruction and Renal Damage in Systemic Disease. J Infect Dis 222, 871–879 (2020).

35. F. Teufel et al., SignalP 6.0 predicts all five types of signal peptides using protein language models. Nat Biotechnol 40, 1023–1025 (2022).

36. C. M. Harding, R. L. Kinsella, L. D. Palmer, E. P. Skaar, M. F. Feldman, Medically Relevant Acinetobacter Species Require a Type II Secretion System and Specific Membrane-Associated Chaperones for the Export of Multiple Substrates and Full Virulence. PLoS Pathog 12, e1005391 (2016).

37. R. L. Kinsella et al., Defining the interaction of the protease CpaA with its type II secretion chaperone CpaB and its contribution to virulence in Acinetobacter species. J Biol Chem 292, 19628–19638 (2017).

38. D. V. Urusova et al., The structure of Acinetobacter-secreted protease CpaA complexed with its chaperone CpaB reveals a novel mode of a T2SS chaperone-substrate interaction. J Biol Chem 294, 13344–13354 (2019).

39. M. F. Haurat et al., The Glycoprotease CpaA Secreted by Medically Relevant Acinetobacter Species Targets Multiple O-Linked Host Glycoproteins. mBio 11 (2020).

40. K. M. Blair et al., Acinetobacter Baumannii Secreted Protease CpaA Inhibits Factor XII-Mediated Bradykinin Generation and Neutrophil Activation. Circ Res 10.1161/CIRCRESAHA.124.324764 (2025).

41. L. Capodimonte et al., OXA beta-lactamases from Acinetobacter spp. are membrane bound and secreted into outer membrane vesicles. mBio 16, e0334324 (2025).

42. K. Masuda, S. Matsuyama, H. Tokuda, Elucidation of the function of lipoprotein-sorting signals that determine membrane localization. Proc Natl Acad Sci U S A 99, 7390–7395 (2002).

43. F. Lauber, G. R. Cornelis, F. Renzi, Identification of a New Lipoprotein Export Signal in Gram-Negative Bacteria. mBio 7 (2016).

44. E. Valguarnera, N. E. Scott, P. Azimzadeh, M. F. Feldman, Surface Exposure and Packing of Lipoproteins into Outer Membrane Vesicles Are Coupled Processes in Bacteroides. mSphere 3 (2018).

45. G. E. Crooks, G. Hon, J. M. Chandonia, S. E. Brenner, WebLogo: a sequence logo generator. Genome Res 14, 1188–1190 (2004).

46. P. Horton et al., WoLF PSORT: protein localization predictor. Nucleic Acids Res 35, W585–587 (2007).

47. J. G. Sklar et al., Lipoprotein SmpA is a component of the YaeT complex that assembles outer membrane proteins in Escherichia coli. Proc Natl Acad Sci U S A 104, 6400–6405 (2007).

48. K. J. Huang et al., Deletion of a previously uncharacterized lipoprotein lirL confers resistance to an inhibitor of type II signal peptidase in Acinetobacter baumannii. Proc Natl Acad Sci U S A 119, e2123117119 (2022).

49. W. Bei et al., Cryo-EM structures of LolCDE reveal the molecular mechanism of bacterial lipoprotein sorting in Escherichia coli. PLoS Biol 20, e3001823 (2022).

50. M. N. Burtnick, P. J. Brett, D. DeShazer, Proteomic analysis of the Burkholderia pseudomallei type II secretome reveals hydrolytic enzymes, novel proteins, and the deubiquitinase TssM. Infect Immun 82, 3214–3226 (2014).

51. V. S. Somvanshi et al., The type 2 secretion Pseudopilin, gspJ, is required for multihost pathogenicity of Burkholderia cenocepacia AU1054. Infect Immun 78, 4110–4121 (2010).

52. H. Zhang et al., Quantitative analysis of cell surface membrane proteins using membrane-impermeable chemical probe coupled with 18O labeling. J Proteome Res 9, 2160–2169 (2010).

53. L. A. Philipp et al., Identification of factors limiting the efficiency of transplanting extracellular electron transfer chains in Escherichia coli. Appl Environ Microbiol 91, e0068525 (2025).

54. C. d’Enfert, A. Ryter, A. P. Pugsley, Cloning and expression in Escherichia coli of the Klebsiella pneumoniae genes for production, surface localization and secretion of the lipoprotein pullulanase. EMBO J 6, 3531–3538 (1987).

55. C. L. Holmes et al., Klebsiella pneumoniae causes bacteremia using factors that mediate tissue-specific fitness and resistance to oxidative stress. PLoS Pathog 19, e1011233 (2023).

56. C. Pineau et al., Substrate recognition by the bacterial type II secretion system: more than a simple interaction. Mol Microbiol 94, 126–140 (2014).

57. M. Gerard-Vincent et al., Identification of XcpP domains that confer functionality and specificity to the Pseudomonas aeruginosa type II secretion apparatus. Mol Microbiol 44, 1651–1665 (2002).

58. A. Shannon et al., The PDZ domain of EpsC is required for extracellular secretion of VesB by the Type II secretion system in Vibrio cholerae. J Bacteriol 10.1128/jb.00144-25, e0014425 (2025).

59. J. M. Gennity, M. Inouye, The protein sequence responsible for lipoprotein membrane localization in Escherichia coli exhibits remarkable specificity. J Biol Chem 266, 16458–16464 (1991).

60. S. Lewenza, M. M. Mhlanga, A. P. Pugsley, Novel inner membrane retention signals in Pseudomonas aeruginosa lipoproteins. J Bacteriol 190, 6119–6125 (2008).

61. S. Y. Tanaka, S. Narita, H. Tokuda, Characterization of the Pseudomonas aeruginosa Lol system as a lipoprotein sorting mechanism. J Biol Chem 282, 13379–13384 (2007).

62. M. G. Kornacker, D. Faucher, A. P. Pugsley, Outer membrane translocation of the extracellular enzyme pullulanase in Escherichia coli K12 does not require a fatty acylated N-terminal cysteine. J Biol Chem 266, 13842–13848 (1991).

63. O. Francetic, A. P. Pugsley, Towards the identification of type II secretion signals in a nonacylated variant of pullulanase from Klebsiella oxytoca. J Bacteriol 187, 7045–7055 (2005).

64. C. D. Jackson-Litteken et al., A chronic Acinetobacter baumannii pneumonia model to study long-term virulence factors, antibiotic treatments, and polymicrobial infections. Nat Commun 16, 7617 (2025).

65. A. E. Sikora, S. R. Lybarger, M. Sandkvist, Compromised outer membrane integrity in Vibrio cholerae Type II secretion mutants. J Bacteriol 189, 8484–8495 (2007).

66. S. Michaelis, C. Chapon, C. D’Enfert, A. P. Pugsley, M. Schwartz, Characterization and expression of the structural gene for pullulanase, a maltose-inducible secreted protein of Klebsiella pneumoniae. J Bacteriol 164, 633–638 (1985).

67. J. Abramson et al., Accurate structure prediction of biomolecular interactions with AlphaFold 3. Nature 630, 493–500 (2024).

