## Supplemental information for "The Type II Secretion System Utilizes AsmA-like Protein GspN to Facilitate Transport of Lipoproteins to the Cell Surface in *Acinetobacter baumannii*"

### Supporting Information

#### The Type II Secretion System Utilizes AsmA-like Protein GspN to Facilitate Transport of Lipoprotein Effector Molecules to the Bacterial Cell Surface

Cameron S. Roberts[1*], Colby Gura[1], Maria Sandkvist[1#]

[1]Department of Microbiology and Immunology, University of Michigan Medical School, Ann Arbor, Michigan, 48109, USA

**This PDF file includes:**

Figures S1 to S10

Tables S1 to S5

SI References

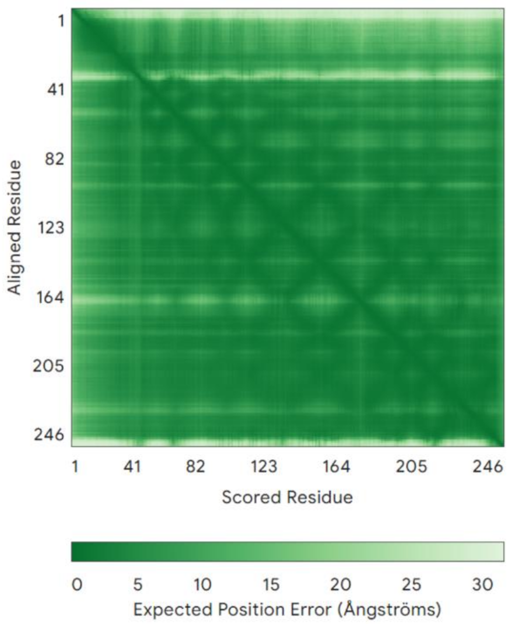


**Supplemental Figure 1. Alignment error for GspN predicted structure.** The primary amino acid sequence of GspN from G414 was used to predict its structure using AlphaFold3 (1). Predicted error shown.

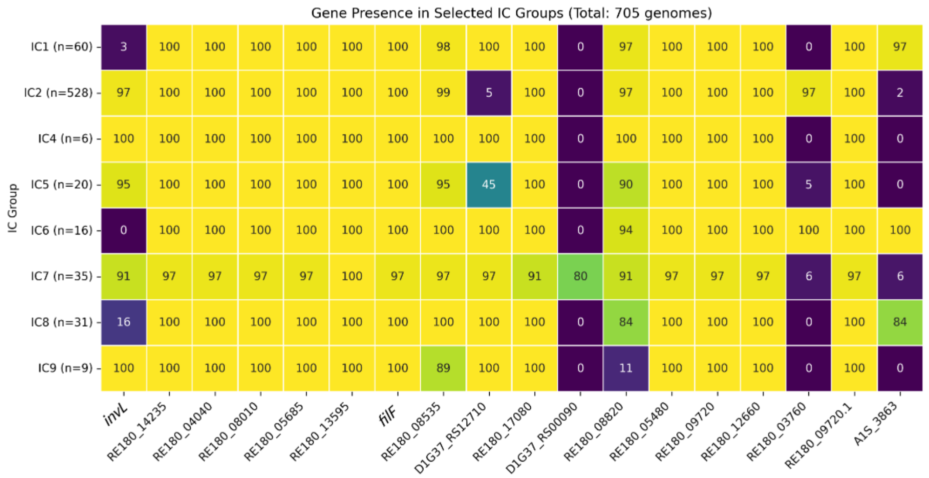


**Supplemental Figure 2. T2SS dependent lipoproteins are highly conserved across IC groups.** Complete genomes were extracted from NCBI and analyzed for presence or absence of individual lipoprotein genes and shown as percent of total within IC groups. Number of genomes analyzed per IC group is listed.

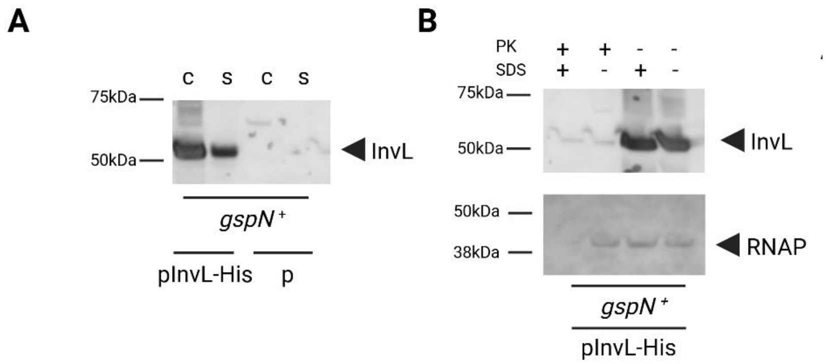


**Supplemental Figure 3. Genomic expression of *gspN* restores InvL secretion and surface localization.** **A.** Overnight cultures from a *gspN* complemented Δ*gspN* strain (*gspN*$^+$) containing empty vector (p) or plasmid expressing InvL with a C-terminal His6 tag were separated into cell (c) and supernatant (s) fractions, run on SDS-PAGE, transferred to a nitrocellulose membrane, and blotted with anti-His6 antibody. Positions of molecular mass markers and InvL are indicated. **B.** Cell fraction from *gspN* complemented strain containing pInvL-His6 was assessed for surface localization of InvL using proteinase K susceptibility with and without cell permeabilization with 1% SDS. For a control, RNAP blots were stained with antibody against RNA polymerase α subunit. Positions of InvL, RNAP and molecular mass markers are shown. Representative western blots are shown (n=3).

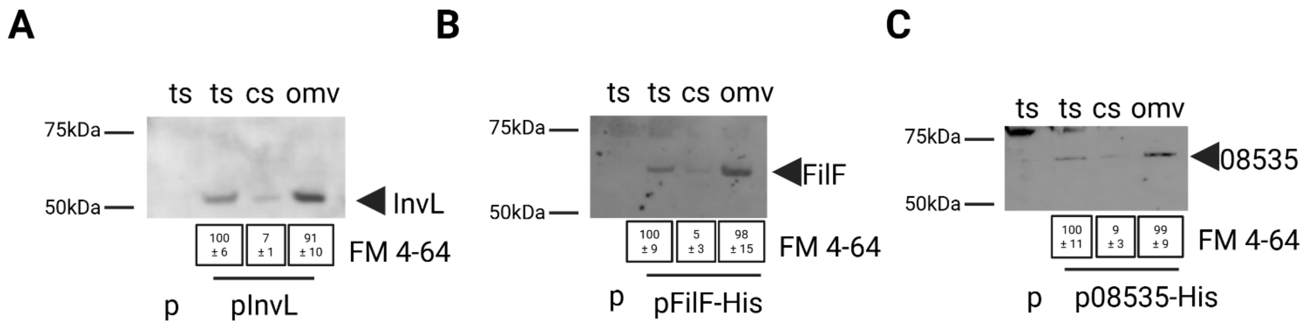


**Supplemental Figure 4. T2SS dependent lipoproteins localize in the pelletable fraction of culture supernatant. A.** Culture supernatants from overnight cultures of WT strain containing empty vector (p) or plasmid expressing InvL with a C-terminal His6 tag were filter sterilized and subjected to high-speed centrifugation to pellet crude outer membrane vesicles (OMVs) and run on SDS-PAGE along with total supernatants (ts) and cleared supernatants (cs), transferred to a nitrocellulose membrane and blotted with His6 antibody. As a control, the fluorescence intensity of samples incubated with FM 4-64 was measured to assess lipid content and indicated below the blot as a mean percentage +/- S. D. of ts fluorescence intensity. **B.** Culture supernatants from overnight cultures of WT strain containing empty vector (p) or plasmid coding for FilF were processed and analyzed as in **A**. **C.** Culture supernatants from overnight cultures of WT strain containing empty vector (p) or plasmid expressing 08535 were processed and analyzed as in **A**. Representative western blots are shown (n=2).

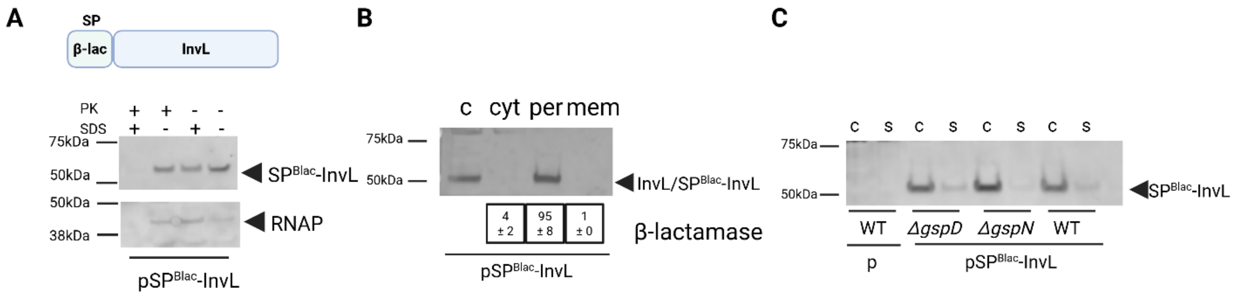


**Supplemental Figure 5. InvL produced with the signal peptide from β-lactamase is primarily localized to the periplasm. A.** Overnight cultures from a WT strain expressing $SP^{\beta lac}$-InvL were assessed for surface localization of $SP^{\beta lac}$-InvL with proteinase K. Positions of molecular mass markers, $SP^{\beta lac}$-InvL, and RNAP are indicated. **B.** Cell fraction (c) from WT cells expressing $SP^{Blac}$-InvL were separated into cytoplasmic (cyt), periplasmic (per), and total membrane (mem) before run on SDS-PAGE, transferred to a nitrocellulose membrane, and blotted with anti-His6. As a control, β-lactamase was assessed with nitrocefin as a marker for periplasmic. Activity is expressed as mean percentage +/- S. D. of the total activity across the three fractions as described in materials and methods. **C.** Overnight cultures from WT and indicated mutant strains containing empty vector (p) or plasmid expressing a chimeric InvL construct containing the signal peptide from β-lactamase ($SP^{\beta lac}$-InvL) as indicated were separated into cell (c) and supernatant (s) fractions, run on SDS-PAGE, transferred to a nitrocellulose membrane, and blotted with anti-His6 antibody. Positions of $SP^{\beta lac}$-InvL and molecular mass markers are indicated. In all cases, representative western blots are shown (n=2).

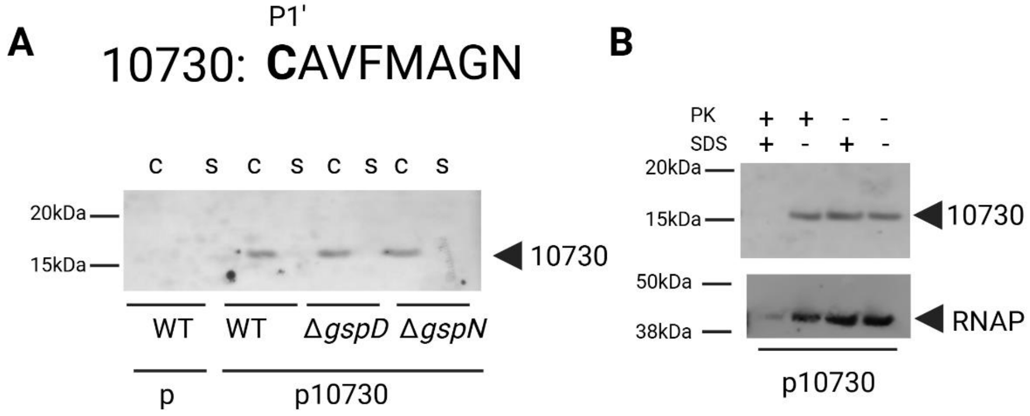


**Supplemental Figure 6. 10730 is not secreted by the T2SS in *A. baumannii*. A.** P1' – P8' residues represented in relation to putative signal peptidase II cleavage site for 10730. Overnight cultures from WT and indicated mutant strains containing empty vector (p) or expressing plasmid-encoded 10730 with C-terminal His6 tag were separated into cell (c) and supernatant (s) fractions, run on SDS-PAGE, transferred to a nitrocellulose membrane, and blotted with anti-His6 antibody. **B.** Cell pellets from overnight cultures from WT and indicated mutant strains containing empty vector (p) or plasmid expressing 10730 with C-terminal His6 tag were assessed for surface localization using proteinase K susceptibility with and without cell permeabilization with 1% SDS. For a control, RNAP blots were re-stained with antibody against RNA polymerase α subunit. 10730 and RNAP are indicated. Representative western blots (n=2).

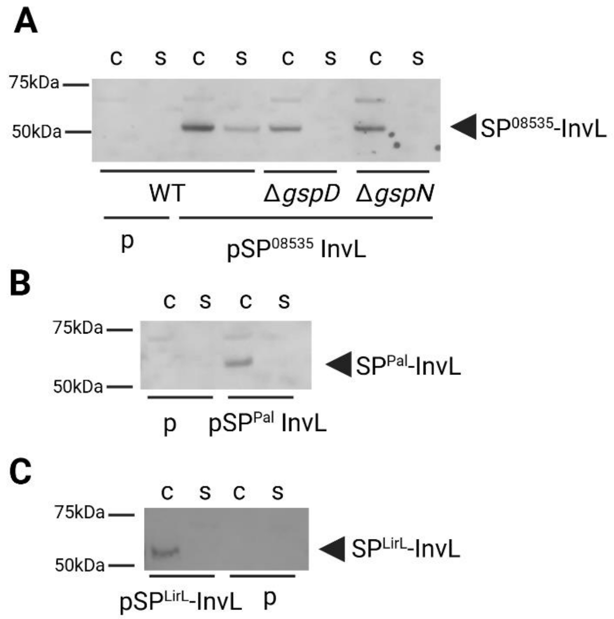


**Supplemental Figure 7. A specific sorting signal is required for secretion of InvL. A.** Overnight cultures from WT and indicated mutant strains containing empty vector (p), or plasmid coding for InvL chimera $SP^{08535}$-InvL with C-terminal His6 tag were separated into cell (c) and supernatant (s) fractions, run on SDS-PAGE, transferred to a nitrocellulose membrane, and blotted with anti-His6 antibody (bottom). Positions of $SP^{08535}$-InvL and molecular mass markers are indicated. **B.** Overnight cultures from WT strain containing empty vector (p) or plasmid coding for the InvL chimera $SP^{Pal}$-InvL with C-terminal His6 tag were separated into cell (c) and supernatant (s) fractions, run on SDS-PAGE, transferred to a nitrocellulose membrane, and blotted with anti-His6 antibody. **C.** Overnight cultures from WT strain containing empty vector (p) or plasmid coding for the InvL chimera $SP^{LirL}$-InvL with C-terminal His6 tag were separated into cell (c) and supernatant (s) fractions, run on SDS-PAGE, transferred to a nitrocellulose membrane, and blotted with anti-His6 antibody. Representative western blots (n=2-3).

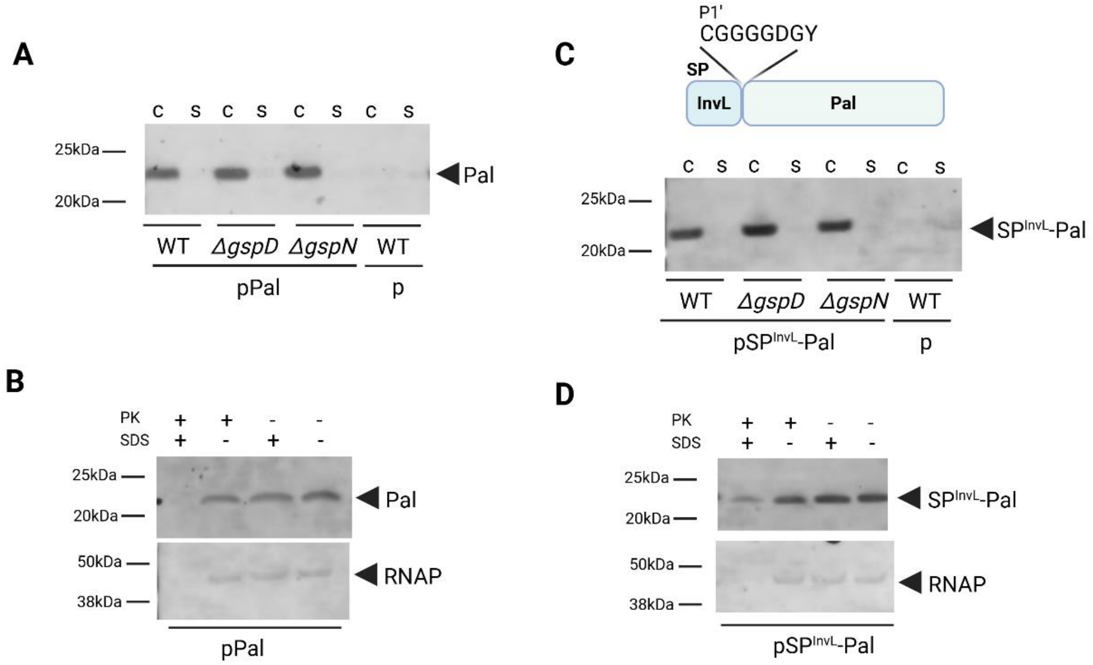


**Supplemental Figure 8. The lipobox from InvL is insufficient to target Pal for secretion via the T2SS. A**. Overnight cultures from WT and indicated mutant strains containing empty vector (p), or plasmid expressing Pal with C-terminal His6 tag were separated into cell (c) and supernatant (s) fractions, run on SDS-PAGE, transferred to a nitrocellulose membrane, and blotted with anti-His6. **B.** Cell pellets from overnight cultures from a WT strain expressing Pal with C-terminal His6 tag were assessed for surface localization using proteinase K susceptibility with and without cell permeabilization with 1% SDS. For a control, RNAP blots of the same samples were stained with antibody against RNA polymerase α subunit. Positions of molecular mass markers, Pal and RNAP are indicated. **C**. Overnight cultures from WT and indicated mutant strains containing empty vector (p), or plasmid expressing chimeric Pal with the signal peptide of InvL ($SP^{InvL}$-Pal) with C-terminal His6 tag were separated into cell (c) and supernatant (s) fractions, run on SDS-PAGE, transferred to a nitrocellulose membrane, and blotted with anti-His6 antibody. **D.** Cell pellets from overnight cultures from a WT strain expressing chimeric Pal with the signal peptide from InvL and a C-terminal His6 tag was assessed for surface localization using proteinase k susceptibility with and without permeabilization with 1% SDS. For a control, RNAP blots were stained with antibody against RNA polymerase α subunit. Positions of molecular mass markers, $SP^{InvL}$-Pal and RNAP are indicated. Representative western blots are shown from at least two blots performed on biological replicas.

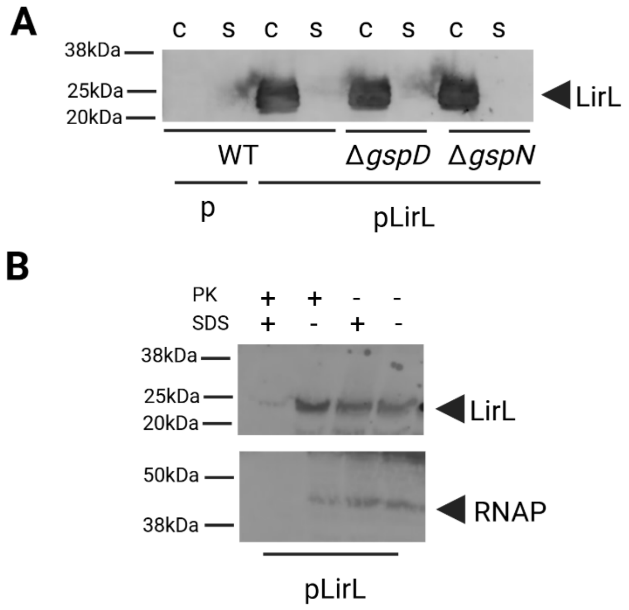


**Supplemental Figure 9. LirL is not secreted or surface displayed. A**. Overnight cultures from WT and indicated mutant strains containing empty vector (p), or plasmid expressing LirL with C-terminal His6 tag were separated into cell (c) and supernatant (s) fractions, run on SDS-PAGE, transferred to a nitrocellulose membrane, and blotted with anti-His6. **B.** Cell pellets from overnight cultures from a WT strain expressing LirL with C-terminal His6 tag were assessed for surface localization using proteinase K susceptibility with and without cell permeabilization with 1% SDS. For a control, RNAP blots of the same samples were stained with antibody against RNA polymerase α subunit. Positions of molecular mass markers, LirL and RNAP are indicated. Representative western blots (n=2).

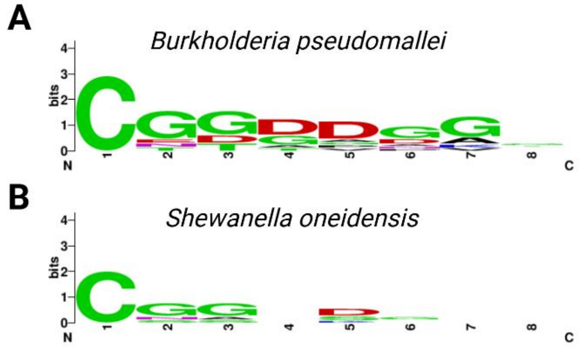


**Supplemental Figure 10. Sorting sequence is conserved across GspN containing T2SS.** Proteomic datasets determining *gspD* dependent secreted proteins from *B. pseudomallei* strain 688 (**A**) and *S. oneidensis* strain MR-1 (**B**) were mined for lipoproteins and sorting motifs were generated for respective strain starting at the P1’ residue relative to the signal peptidase II cleavage site.

**Table S1. Putative Inner membrane proteins from G414.**

| Protein | Sorting Motif |
|---|---|
| RE180_01550 | **C**DSRIDAV |
| RE180_03225 | **C**NQQEVPK |
| RE180_03575 | **C**ILIFSAY |
| RE180_07055 | **C**QVVNVKQ |
| RE180_08740 | **C**DSKEVAQ |
| RE180_10800 | **C**GAVYSAI |
| RE180_10945 | **C**GGDMVLL |
| RE180_11055 | **C**GKQQNEP |
| RE180_11805 | **C**NKQPAQT |
| RE180_16525 | **C**QNMSPSD |
| RE180_16530 | **C**QNASPES |
| RE180_17100 | **C**TSLGPNS |

**Bold** indicates predicted P1' cysteine residue relative to signal peptidase II cleavage site.

**Table S2. Putative Outer membrane proteins from G414.**

| Protein | Sorting Motif |
|---|---|
| RE180_02600 | **C**LSFGPLK |
| RE180_02925 | **C**SPVIFAE |
| RE180_03640 | **C**QSMRGPE |
| RE180_04385 | **C**ASRKPAT |
| RE180_05035 | **C**ASKPQIN |
| RE180_05700 | **C**SLAPEYQ |
| RE180_07535 | **C**SGSIQSS |
| RE180_08655 | **C**WAMYEVS |
| RE180_08730 | **C**VNMQAPQ |
| RE180_11885 | **C**SSTPQSA |
| RE180_12500 | **C**NKHENKT |
| RE180_13675 | **C**SIFGVYK |
| RE180_13725 | **C**QTTGNNL |
| RE180_13950 | **C**SSNPSKK |
| RE180_15305 | **C**AAVVKTP |
| RE180_15475 | **C**QTSQTVK |
| RE180_17745 | **C**AVTSGLQ |

**Bold** indicates predicted P1' cysteine residue relative to signal peptidase II cleavage site.

**Table S3. Putative T2SS dependent lipoproteins from *Burkholderia pseudomallei*.**

| Protein | Sorting Motif |
|---|---|
| BURPS668_A2859 | **C**GGDDGAG |
| BURPS668_3144 | **C**GGDDGGG |
| BURPS668_3846 | **C**GGGGDGG |
| BURPS668_3613 | **C**GGDDGGT |
| BURPS668_3884 | **C**ETAVPAA |
| BURPS668_3613 | **C**GGDDGGT |
| BURPS668_3454 | **C**NGDDAGS |
| BURPS668_A1338 | **C**GDGANGE |
| BURPS668_A2630 | **C**GDGDGVV |
| BURPS668_2270 | **C**TTTPDKP |

**Bold** indicates predicted P1' cysteine residue relative to signal peptidase II cleavage site.

**Table S4. Putative T2SS dependent lipoproteins from *Shewanella oneidensis*.**

| Protein | Sorting Motif |
|---|---|
| SO_1429 | **C**NSGSSDV |
| SO_1778 | **C**GGSDGNN |
| SO_1779 | **C**GGSDGKD |
| SO_0404 | **C**GAGDEPY |
| SO_A0112 | **C**GGDKGFL |
| SO_A0110 | **C**SGESRNN |

**Bold** indicates predicted P1' cysteine residue relative to signal peptidase II cleavage site.

**Table S5. Plasmids and primers used in study.**

| Plasmid/strain | Description | Fwd Primer 5' to 3' | Rev Primer 5' to 3' |
|---|---|---|---|
| G414 | Clinical isolate from University of Michigan | | |
| Δ*gspD* | G414 with markerless, in frame deletion of *gspD* (2) | | |
| Δ*gspN* | G414 with markerless, in frame deletion of *gspN* (2) | | |
| *gspN*+ | Complemented Δ*gspN* strain with *gspN* reintroduced in genome in the intergenic region upstream of the *gspN-gspD* operon | Upstream: gtcgacaaaaagtaaatttatagcgttattc<br>Downstream: ttattacaaggtggtaactaattttatatgaagtgaattggcg | Upstream: ttgcttagactttttcttcataagcaatcgggcttg<br>Downstream: cccgggtttcctaactcttatggtgacg |
| MC1061 | *E. coli* cloning strain (3) | | |
| pRK2013 | Helper Strain, (4) | | |
| SY327λpir | Helper strain (5) | | |
| KPRR1 | (6) | | |
| KPRR1 *pulA*::tn | (6) | | |
| KPRR1 *pulN*::tn | (6) | | |
| pSC-B-Amp/Kan | Cloning vector (Agilent) | | |
| pMMB67 | Expression vector (7) | tggctgtgcaggtcgtaaatcac | tactcaggagagcgttcaccgacaaacaac |
| pCVD442 | Suicide vector (5) | | |
| pInvL | pMMB67, C-terminal 6-his | atcttatacctttaggaaagattgggg | ctgtagttgccgttgtattaccatttgaacagtttggatc |
| pCpaA | (2) | | |
| pBlac | (8) | | |
| pSP$^{βlac}$-InvL | InvL with the SP from β-lactamse | ttttgccttcctgtttttgctggcgggggaggcggtgatg | accgcctcccccgccagcaaaaacaggaaggcaaaatgc |
| pSP$^{LirL}$-InvL | C-terminal 6-his | tgttctaagaaagaagaagctccatattaccaaaatggttcatcaaataac | ATTTGATGAACCATTTTGGTAATAtggagcttcttctttcttagaacaacc |
| pSP$^{Pal}$-InvL | InvL with SP from Pal | aagtcgtaagccagcaggctattaccaaaatggttcatc | accattttggtaatagcctgctggcttacgacttgc |
| pFilF | C-terminal 6-his | tttcggtattacgttaaaaataagtaggg | ttaatggtgatgatggtgatgagacttagttttaattgctttacaagctgg |
| p083535 | C-terminal 6-his | tgcgtattttttatgcatcttaattcatc | ttaatggtgatgatggtgatgtttaatgactttaatacgagtcgtagcc |
| p10730 | C-terminal 6-his | aaattggatatgatgagcgcgtaatactttaagc | ttaatggtgatgatggtgatgttgttcttttacttttggaaaagcttgattaaagtc |
| pLirL | C-terminal 6-his | aatctagagtgtagattaatggtggtgatggtggtg | tctgcagaatctagagtgtagattaatggtggtgatggtggtg |
| pPal | C-terminal 6-his | tttttattccattgatggagatg | ttaatggtgatgatggtgatgttttaatagaggaggaaccgcttc |
| pSP$^{InvL}$-Pal | Pal with SP from InvL | gggaggcggtgatacaacggcaactacaggtac | ctgtagttgccgttgtatcaccgcctccccc |
| pPulN | pMMB67, from KPRR1 | atctgctgctggagcg | gataacgcggctagtgtc |
| pGspN | (2) | | |

**Supplemental References**


1. J. Abramson *et al.*, Accurate structure prediction of biomolecular interactions with AlphaFold 3. *Nature* **630**, 493-500 (2024).
2. T. L. Johnson, U. Waack, S. Smith, H. Mobley, M. Sandkvist, Acinetobacter baumannii Is Dependent on the Type II Secretion System and Its Substrate LipA for Lipid Utilization and In Vivo Fitness. *J Bacteriol* **198**, 711-719 (2015).
3. C. T. Lutz, W. C. Hollifield, B. Seed, J. M. Davie, H. V. Huang, Syrinx 2A: an improved lambda phage vector designed for screening DNA libraries by recombination in vivo. *Proc Natl Acad Sci U S A* **84**, 4379-4383 (1987).
4. V. C. Knauf, E. W. Nester, Wide host range cloning vectors: a cosmid clone bank of an Agrobacterium Ti plasmid. *Plasmid* **8**, 45-54 (1982).
5. M. S. Donnenberg, J. B. Kaper, Construction of an eae deletion mutant of enteropathogenic Escherichia coli by using a positive-selection suicide vector. *Infect Immun* **59**, 4310-4317 (1991).
6. C. L. Holmes *et al.*, Klebsiella pneumoniae causes bacteremia using factors that mediate tissue-specific fitness and resistance to oxidative stress. *PLoS Pathog* **19**, e1011233 (2023).
7. J. P. Furste *et al.*, Molecular cloning of the plasmid RP4 primase region in a multi-host-range tacP expression vector. *Gene* **48**, 119-131 (1986).
8. A. Shannon *et al.*, The PDZ domain of EpsC is required for extracellular secretion of VesB by the Type II secretion system in Vibrio cholerae. *J Bacteriol* 10.1128/jb.00144-25, e0014425 (2025).
